# Simulations of the infant gut microbiota suggest that complex ecological interactions regulate effects of human milk oligosaccharides on microbial mucin consumption

**DOI:** 10.1101/2024.07.15.603541

**Authors:** David M. Versluis, Clair Wijtkamp, Ellen Looijesteijn, Jan M. W. Geurts, Roeland M. H. Merks

## Abstract

Intestinal mucin acts as a barrier protecting the infant gut wall against diseases such as colitis and rotavirus. *In vitro* experiments have shown that the gut microbiota of breastfed infants consumes less mucin than the microbiota of non-breastfed infants, but the mechanisms are incompletely understood. The main difference between human milk and most infant formulas is the presence of human milk oligosaccharides (HMOs) in human milk. We hypothesize that HMOs protect mucin by stimulating non-mucin consuming bacteria. To understand the underlying mechanisms we developed a computational model that describes the metabolism and ecology of the infant gut microbiota. Model simulations suggest that extracellular digestion of the HMO 2’-fucosyllactose by the mucin-consumer *Bifidobacterium bifidum* may make this species vulnerable to competitors. The digestion products of HMOs become ‘public goods’ that can be consumed by competing species such as *Bacteroides vulgatus* instead.*Bifidobacterium longum*, which does not consume mucin or produce public goods, can then become dominant, despite growing less efficiently on HMOs in monocultures than *B. bifidum*. In conclusion, our model simulations suggest that, through complex ecological interactions, HMOs may help lower mucin consumption by stimulating the non-mucin consumer *B. longum* at the expense of the mucin consumer *B. bifidum*.

## 1. Introduction

The interior of the gastrointestinal tract is covered by a layer of mucin which protects the gut against diseases such as colitis [1] and rotavirus infection [2]. The mucin layer consists largely of mucin glycoproteins. Some species of intestinal bacteria, such as *Bacteroides* spp. and *Bifidobacterium bifidum*, can consume these mucins [3,4]. As mucin protects the gut against infection, consumption of intestinal mucins by the residing microbiota potentially increases the risk of disease [5]. It is thought that breastfeeding shapes the infant gut microbiota in such a way that the microbiota consumes less mucin [6]. This hypothesis is based on *in vitro* observations showing that microbiota from the feces of breastfed infants consume mucins more slowly than the microbiota from the feces of non-breastfed infants [6].

As infant formula at the time of this study did not contain any human milk oligosaccharides (HMO), we hypothesize that it is the HMO in human milk that shaped the microbiota of the breastfed infants to consume less mucin. HMOs are the most abundant component of human milk after lactose and lipids [7]. They are not digested by the host, but exclusively by intestinal bacteria, thereby shaping the microbiota [8]. About 200 HMO structures exist, each of which consists of a core of lactose with other sugars attached to it [9]. We will focus in particular on the potential effects of the HMO 2’-fucosyllactose (2’-FL), as it is the most abundant HMO in most human milk [10] and its digestion by the infant gut microbiota is well-characterized [11]. We also examine galacto-oligosaccharides (GOS), which have a similar structure and are consumed by the same bacteria, but are not HMOs [12]. GOS are frequently added to commercial infant formula [13]. HMOs may decrease mucin consumption by stimulating non-mucin consumers at the expense of mucin consumers. Indeed, the non-mucin consumer *Bifidobacterium longum* specializes in HMOs, including 2’-FL [14], and a high abundance of *B. longum* in the infant gut microbiota lowers total mucin consumption in breastfed infants [15]. Paradoxically, in *in vitro* experiments, the mucin consumer *Bifidobacterium bifidum* digests and takes up HMOs such as 2’-FL more efficiently than *B. longum* [11,16], but *B. bifidum* is much less abundant than *B. longum* in the infant gut [17,18]. We hypothesize that differences in HMO digestion strategies cause *B. longum* to be more abundant than *B. bifidum in vivo*.

The carbohydrate metabolism of *B. bifidum* and *B. longum* is similar, as both species use bifid shunt metabolism to metabolise sugars [19], but the species crucially differ in how they break down and transport HMOs into the cell [11]. *B. longum* strains that consume HMO, particularly *B. longum* ssp. *infantis*, import most HMOs, including 2’-FL, into the cell using ATP-dependent active transport [8]. In contrast, *B. bifidum* digests most HMOs, including 2’-FL, outside the cell, secreting enzymes that break down HMOs into smaller sugars such as lactose [7]. The import of lactose produced from 2’-FL does not require active transport [20]. Thus, *B. bifidum*’s metabolism of 2’-FL is thought to be more efficient. However, a major disadvantage of extracellular digestion is that other species can take up the digestion products [21]. We hypothesize that cross-feeding by other species on the digestion products of *B. bifidum* explains the relatively low abundance of *B. bifidum in vivo*.

Extracellular products that can benefit other species instead of the producer are a type of public goods [22]. The role of public goods in inter-species interaction has been studied in detail in many systems, in particular in budding yeast (*Saccharomyces cerevisiae*) [23]. Yeast cells can secrete the enzyme invertase that converts sucrose into fructose and glucose. With this enzyme yeast can grow in glucose-limited medium rich in sucrose. However, the cell that secretes invertase can only take up around 1% of the glucose produced, as the rest is lost to the environment. This way, glucose functions as a public good. Because the production of invertase is costly, it can be advantageous for a yeast cell to not produce invertase, but only take up glucose produced by invertase from other cells. In yeast communities invertase producers and invertase non-producers typically co-exist. Co-existence is possible because they are playing a ‘snowdrift game’, in which whatever strategy is less common (producing or not producing invertase) has an advantage over the more common strategy [23]. The extracellular digestion of oligosaccharides by *B. bifidum* has also been considered as a form of public goods production [21], and may be influenced by the same mechanisms. In our model of the infant microbiota, *B. bifidum* is a producer of public goods from 2’-FL and every other species is a non-producer. Strong pressure from cross-feeding or ‘stealing’ by the surrounding bacterial populations may then explain why *B. bifidum* is less abundant than other bacteria *in vivo*. As in yeast, an advantage of the less common strategy (producing) over the more common strategy (non-producing) may explain why *B. bifidum* is often present at low abundance in the infant gut.

Concretely, we investigated (1) whether 2’-FL in milk can explain the reduced mucin consumption observed in the microbiota of breastfed infants, and (2) whether public goods metabolism of 2’-FL can explain the relatively low abundance of the mucin consumer *B. bifidum* compared to *B. longum* in the infant gut microbiota. As these mechanisms involve an interplay between the molecular, population and physical levels, we used a multiscale mathematical model that uses flux balance analysis (FBA) on genome-scale metabolic models (GEMs). Such models have previously been used to model dynamic bacterial interactions with spatial separation, such as in the gut microbiota [24–27]. We based the model on our previous microbiota models [26,28,29], which modelled microbial interactions in the adult and infant gut microbiota, including relevant host factors such as flow rate and the presence of initial oxy-gen. These models have previously been used to generate predictions on the effects of factors such as diarrhoea, oxygenation, and HMO supplementation on microbiota composition and metabolism [26,28,29]. We have extended these models to also include species-specific public goods-producing and non-public goods-producing metabolism of mucin and 2’-FL.

Briefly, we show that our model can reproduce cross-feeding between *B. bifidum* and *Anaerobutyricum hallii* as observed *in vitro* and that it predicts that:

1. 2’-FL reduces mucin consumption by causing a higher abundance of *B. longum*.
2. The low abundance of *B. bifidum* can be explained by other species consuming the public goods it produces from 2’-FL.
3. *B. longum* is more abundant than mucin consumers because its 2’-FL metabolism is intracellular, and so does not produce public goods.

## 2. Results

### 2.1. Public goods model reproduces in vitro cross-feeding between mucin consuming and non-consuming species

We first tested if our model system could reproduce consumption of mucin by a cross-feeding community of two species. *In vitro*, *Anaerobutyricum hallii* cannot grow on mucin in single culture, whereas it can grow on mucin in co-culture with *B. bifidum* [30]. *B. bifidum* digests mucin extracellularly, which leads to cross-feeding metabolites becoming available in the medium as public goods [30]. *A. hallii* can then grow on these cross-feeding metabolites [30]. We first investigated whether our computer model could reproduce this cross-feeding.

We simulated this two-species system by extending a multiscale model as previously described [28,29]. Briefly, the model integrates simulations of the metabolism of bacterial species to generate predictions for microbial abundances and metabolic interactions over space and time. Bacterial metabolism is predicted using FBA on GEMs. The growth rates of the bacteria are given by a biomass reaction whose rate is maximized using FBA. The default biomass reaction in the model uses ATP production as a proxy for biomass production. We have extended this model with species-specific mucin metabolism and extracellular public goods production from mucin and 2’-FL. The extracellular public goods production is separate from the FBA predictions of metabolism (Fig. 1A&B). We distributed all nutrients and metabolites uniformly across all lattice sites at the start of each timestep. We supplied the system only with water and mucin, and only used GEMs of *B. bifidum* and *A. hallii*. We used a fucosylated core-2 mucin structure to represent all mucin oligosaccharides. For details see the methods sections ‘Non-spatial model’ and ‘Nutrient input’.

**Figure 1.**
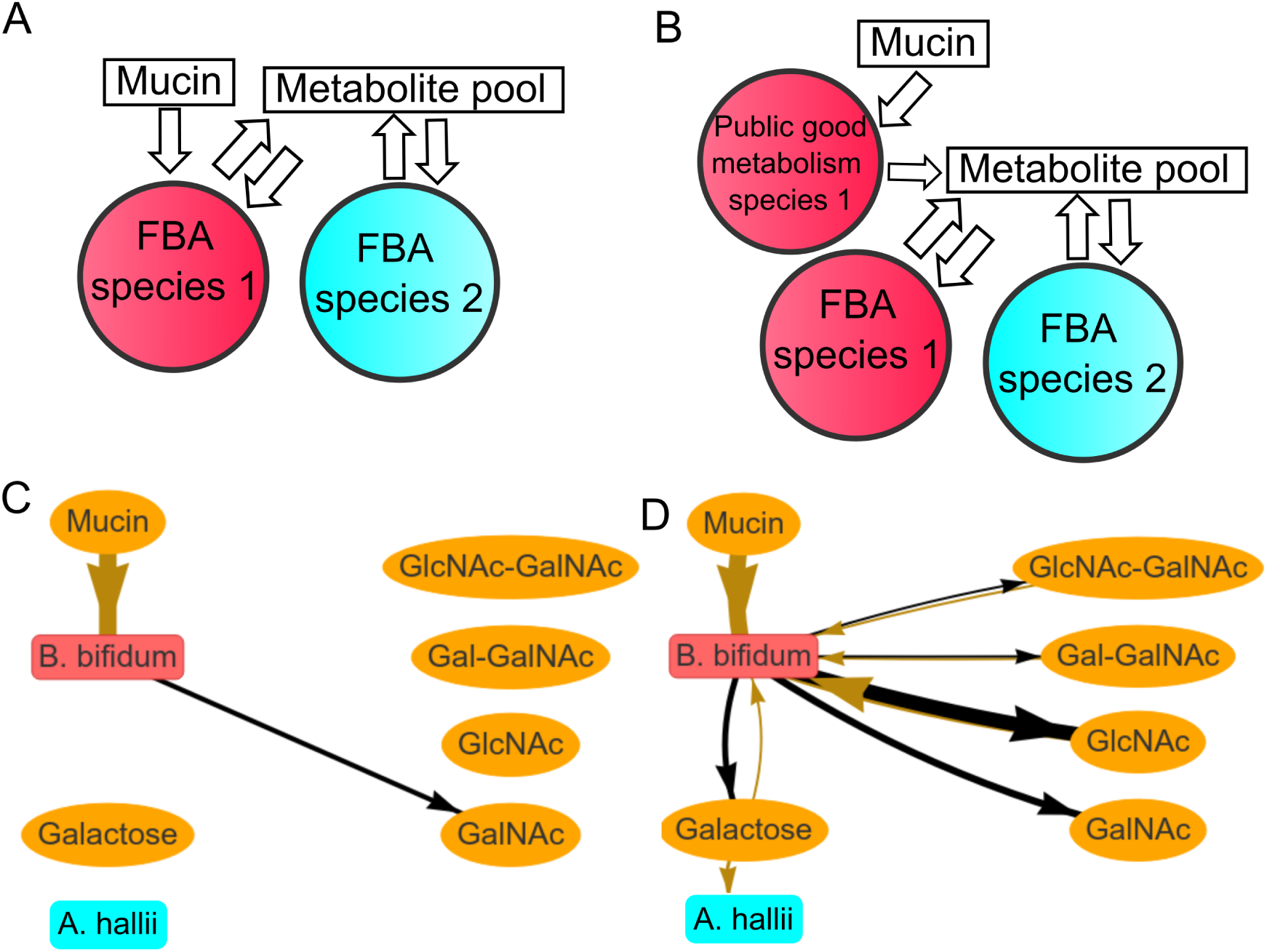
Model outline and effect of the public goods model. (A) Schematic of the non-public goods model without spatial separation (see methods section ‘Non-spatial model’) fed with mucin. Mucin is consumed by specific species, and all bacterial populations share a common nutrient pool. (B) Schematic of the public goods model without spatial separation fed with mucin. Mucin is consumed separately from other substrates to represent extracellular metabolism, and all bacteria need to retrieve the metabolites of mucin metabolism from the common pool. (C,D) Metabolic network for *B. bifidum* and *A. hallii* in a single simulation with no spatial separation and mucin as the only nutrient source not using the public goods model (C) and using the public goods model (D). Yellow arrows indicate uptake, black arrows indicate production. Line width is scaled with the flux per metabolite, multiplied by the carbon content of the molecule, with a minimum threshold of 100 µmol atomic carbon.

We compared two model variants that attempt to capture the same biological processes with different representations. In the first model variant, the ‘control model’, we assumed that bacterial populations have access to all metabolites they produce, even when they are produced extracellularly. We implement this by letting all reactions be represented by FBA. In this model the FBA solution selects what digestion products of mucin to create, and these were first available in the local lattice site, such that species that produced the digestion enzymes had exclusive benefits from the digestion products in the timestep when they were created. In a second variant, the ‘public goods model’, we assumed that the products of extracellular metabolism become available to other bacterial populations when they are produced. We implement this with a separate handling of public goods production from mucin and HMO. In this model, all possible mucin digestion products were produced outside of the FBA, and these diffused before they became available for uptake by the populations. This potentially made these products available to populations at other sites.

In simulations of the control model, *A. hallii* did not grow, even when *B. bifidum* produced cross-feeding products such as GalNAc (Fig. 1C). However, in the ‘public goods’ model variant, the model predicted that *B. bifidum* also released galactose into the medium. *A. hallii* could then grow in co-culture with *B. bifidum* by consuming the galactose (Fig. 1D), in agreement with experimental observations [30].

Summarizing, the multiscale model correctly reproduced the cross-feeding of *A. hallii* and *B. bifidum*. Therefore the model with public goods metabolism of mucin provides a good basis for simulating more complex situations, as we will show in the next sections.

### 2.2. Public goods model reproduces in vivo microbiota composition and mucin consumption

We next turned our attention to mucin consumption in the context of a more diverse infant gut microbiota. Two observations suggest that breastfeeding shapes the infant microbiota such that mucin consumption is reduced. Firstly, *in vitro* data suggest that the microbiota from breastfed infants consume less mucin than the microbiota from non-breastfed infants [6]. Secondly, *in vivo*, a breastfed infant microbiota dominated by *B. longum* consumes less mucin [15].

We tested if we could reproduce and analyse these observations in a model of a complex microbiota, as described by the full infant gut microbiota model as introduced previously [28,29]. We extended this microbiota consortium with microbial reactions for mucin production and consumption, and extracellular digestion of the HMO 2’-FL (see methods section ‘Changes to genome-scale metabolic models’). Briefly, the spatial model represents the infant colon with a regular square lattice of 225 *×* 8 boxes of 2 *×* 2 mm, (Fig. 2 A&B). Nutrients are introduced into the system at regular intervals, and populations, nutrients, and metabolites diffuse through the system. Nutrients and metabolites advect through the system, and are removed at the distal end. Mucin is produced at the upper boundaries of the system, to representing mucin production by the intestinal wall. The lower boundary represents the center of the intestinal lumen, the left boundary represents the entrance of the intestine, and the right boundary represents its exit.

**Figure 2.**
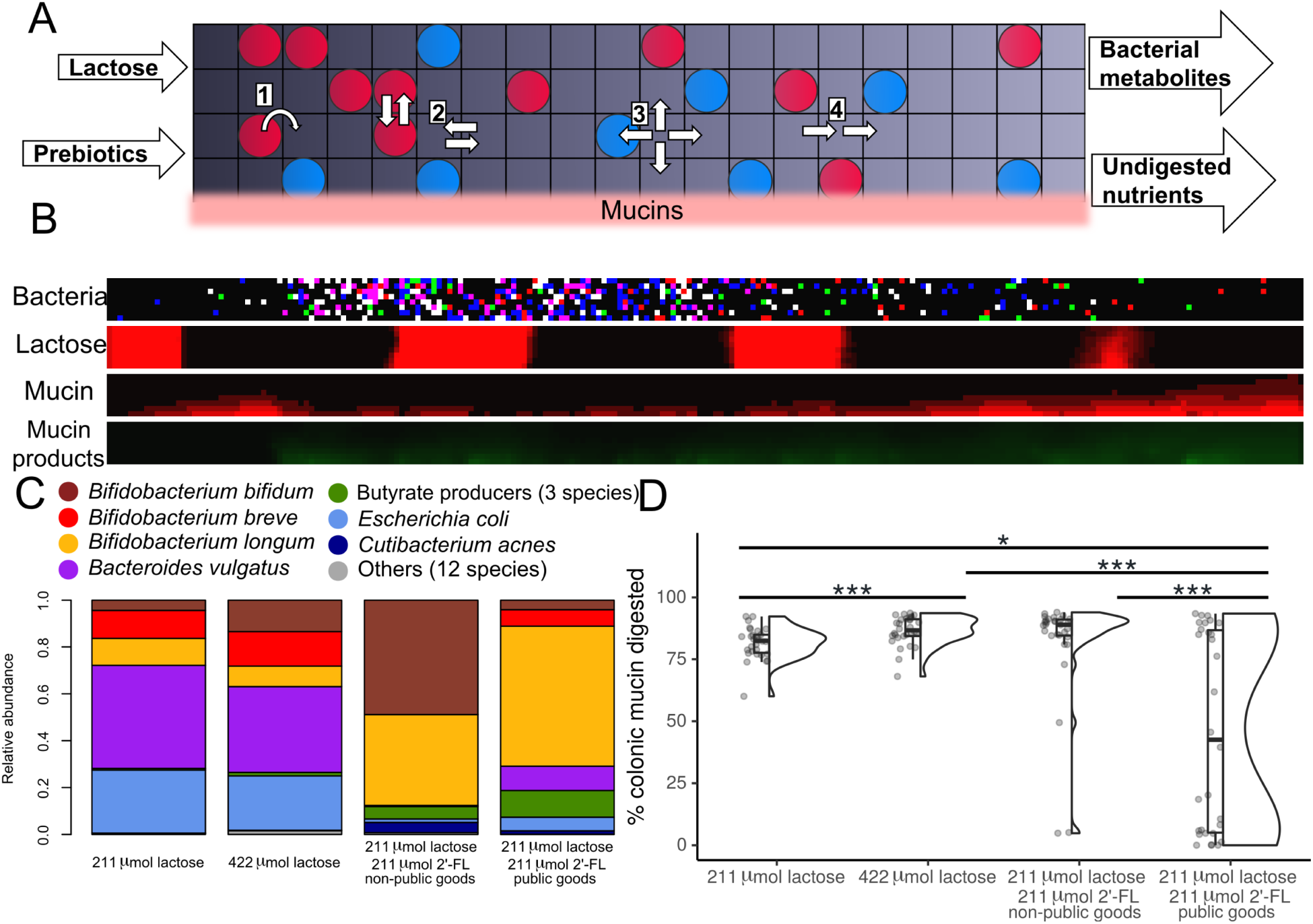
2’-FL metabolism stimulates a microbiota that consumes less mucin and has dominant *B. longum* exclusively in the public goods model. (A) Schematic of the model. Circles represent populations, color represents species. Lactose and 2’-FL enter left (proximally). Mucin enters at the bottom. All nutrients and metabolites move distally (right) and leave the system there. Numbers indicate processes in the model: (1) Metabolism, growth, and division of populations (2) Diffusion of populations (3) Diffusion of metabolites (4) Advection of metabolites (B) Screenshot of a model simulation, showing from top to bottom the bacterial layer, lactose, mucin, and mucin products. Color indicates species in the bacterial layer. Brightness indicates growth in the bacterial layer, and concentration in the other layers. (C) Average relative abundance of bacterial species at the end of 21 days with four different conditions: (1) with 211 µmol lactose, (2) 422 µmol lactose, (3) 211 µmol lactose and 211 µmol 2’-FL without public goods 2’-FL metabolism, or 211 µmol lactose and (4) 211 µmol 2’-FL with public goods 2’-FL metabolism. n=30 for each condition. (D) Amount of colonic mucin digested by the microbiota as a percentage of total mucin released into the gut over the final three hours of the model, per condition of C. n=30 for each condition NS: Not significant, *: p<0.05, **:p<0.01, ***:p<0.001

The model contains 21 bacterial species, each represented by a GEM (Table 1). The species were selected based on *in vivo* data [17], as described in methods section ‘Species composition’. Of these species, only *B. bifidum* and *B. longum* could digest 2’-FL in the model. Again we simulated a public goods model variant alongside a non-public goods model variant as control. *B. bifidum* produced public goods from 2’-FL only in the public goods model, but *B. longum* always digested 2’-FL without producing public goods, even in the public goods model of 2’-FL metabolism. *B. bifidum*, *Bacteroides vulgatus*, and *Parabacteroides distasonis* could digest mucin, all of which produced public goods.

**Table 1.** Species and subspecies included in the computational model. Color indicates color used in figures.

| Name | Phylum | Mucin consumption | 2'-FL consumption |
| --- | --- | --- | --- |
| *Bifidobacterium longum* ssp. *infantis* | Actinomycetota | No | Without public goods |
| *Bifidobacterium breve* | Actinomycetota | No | No |
| *Bifidobacterium bifidum* | Actinomycetota | With public goods | With public goods |
| *Collinsella aerofaciens* | Actinomycetota | No | No |
| *Cutibacterium acnes* | Actinomycetota | No | No |
| *Rothia mucilaginosa* | Actinomycetota | No | No |
| *Eggerthella* sp. YY7918 | Actinomycetota | No | No |
| *Streptococcus oralis* | Bacillota | No | No |
| *Staphylococcus epidermidis* | Bacillota | No | No |
| *Gemella morbillorum* | Bacillota | No | No |
| *Enterococcus faecalis* | Bacillota | No | No |
| *Lactobacillus gasseri* | Bacillota | No | No |
| *Ruminococcus gnavus* | Bacillota | No | No |
| *Veillonella dispar* | Bacillota | No | No |
| *Anaerobutyricum hallii* | Bacillota | No | No |
| *Roseburia inulinivorans* | Bacillota | No | No |
| *Clostridium butyricum* | Bacillota | No | No |
| *Parabacteroides distasonis* | Bacteroidota | With public goods | No |
| *Bacteroides vulgatus* | Bacteroidota | With public goods | No |
| *Haemophilus parainfluenzae* | Pseudomonadota | No | No |
| *Escherichia coli* SE11 | Pseudomonadota | No | No |

We simulated the model with four conditions, differing in nutrient input and handling of public goods 2’-FL metabolism: (1) a nutrient input of 211 µmol of lactose per 60 timesteps (3 hours) (2) a nutrient input of 422 µmol of lactose per 60 timesteps (3) a nutrient input of 211 µmol of lactose and 211 µmol of 2’-FL per 60 timesteps, without public goods 2’-FL metabolism, and (4) a nutrient input 211 µmol of lactose and 211 µmol of 2’-FL per 60 timesteps, with public goods 2’-FL metabolism. We did this to examine which approach better reproduced *in vivo* observations. 211 µmol of lactose per 60 timesteps is our best estimate for lactose input into the infant gut, but we also included an increased amount of lactose in condition 2 to control for the increased amount of sugars present in the system when 2’-FL is included. This allowed us to distinguish between an effect of 2’-FL and an effect of total sugar input.

In the conditions with 2’-FL (1 and 2), the simulations predicted a high abundance of all *Bifidobacterium* species, *Escherichia coli*, and *Bacteroides vulgatus*, (Fig. 2C) and a high mucin consumption (Fig. 2D). The model predictions without 2’-FL matched well with the outcomes commonly observed in non-breastfed infants, who generally did not consume HMO. As observed in the formula-fed infant gut microbiota, the model predicts a diverse microbiota, with *Bifidobacterium*, *Bacteroides* and *Escherichia* as major groups [17,31,32] and high mucin consumption [6].

In the condition with 2’-FL and without public goods 2’-FL metabolism, the model predicted that *B. longum* and *B. bifidum* became highly abundant, leading to a high mucin consumption (Fig. 2C&D). In the condition with 2’-FL and public goods 2’-FL metabolism, we observed a dominance of *B. longum* and lower mucin consumption (Fig. 2C&D). Mucin consumption was not lower in all simulations, as the distribution was bimodal (Fig. 2D). Moreover, the public goods model outcomes matched the *in vivo* data on breastfed infants better than the non-public goods model, as *B. longum* is generally more abundant than *B. bifidum* in the infant gut microbiota [17,18], and *in vitro* mucin consumption by the microbiota of breastfed infants is generally lower [6]. A high *B. longum* abundance is also associated with lower mucin consumption in breastfed infants [15].

We concluded that predictions of the public goods model of 2’-FL metabolism matched observations of the inhibitory effect of breastfeeding and *B. longum* abundance on mucin consumption better than the non-public goods model. We hypothesise that this happens because 2’-FL stimulates *B. longum*, and that *B. longum* can suppress the mucin consumer *B. bifidum* through complex ecological interactions. We will further examine this in section ‘*B. bifidum* produces cross-feeding metabolites that may allow it to be exploited by other bacteria’.

### 2.3. Model outcomes robust for different oligosaccharides but not for all parameters or growth assumptions

We next analysed the robustness of our results by performing a parameter sensitivity analysis. We analyzed 21 different parameter changes in total. We have collected these changes and the major outcomes in table 2. In the first 15 of the parameter changes we only changed the value of existing parameters. In the following 6 we also explored other relevant conditions, which we will now explain in some more detail. Firstly, we investigated the effects of replacing 2’-FL with GOS, a mixture of oligosaccharides that is commonly used in infant nutrition. To model GOS we implemented alternative digestion reactions in the GEMS. Notably, all *Bifidobacterium* species break down chains of four or more GOS sugars extracellularly [33–36]. Chains of three sugars are imported using active transport by *B. longum* and *Bifidobacterium breve*, but broken down extracellularly by *B. bifidum*. A full description of the digestion of GOS in the model is in methods section ‘Changes to genome-scale metabolic models’. The second major change we investigated was a different mucin structure. In all simulations so far we considered a fucosylated core-2 mucin structure, but due to genetic variation, some humans have intestinal mucins that are not fucosylated [37,38]. To examine the consistency of our results for the non-fucosylated core-2 mucin we repeated the simulations, including the simulations with GOS, with a non-fucosylated core-2 mucin. A detailed description is in methods section ‘Changes to genome-scale metabolic models’. Finally, to assess the effects of the simplifications we made in modelling biomass, we next tested four biomass proxies that different from our earlier assumption that ATP production rate determines biomass production: (1) ATP production rate and acetate production rate, (2) only acetate production rate, (3) ATP production rate and pyruvate production rate, or (4) ATP production rate and ketoglutarate production rate. Acetate, pyruvate, and ketoglutarate are used as building blocks for bacterial biomass in *E. coli in vitro* [39]. Alternative biomass reactions 1,3, and 4 required respectively 1, 3, and 5 mol of ATP to be produced per mol of carbohydrate. This brings the ratio between carbon atoms and ATP in line with those in the default biomass reactions of the GEMs we used [40].

**Table 2.** Parameter changes and outcomes.

| Parameter changed | Magnitude | Figure | Major differences in outcomes |
| --- | --- | --- | --- |
| Public goods production | $\times 0.1$ | S1 | - |
| | $\times 0.5$ | S1 | - |
| | $\times 2$ | S1 | - |
| Enzymatic constraint | $\times 0.5$ | S2A | Butyrate producers extinct with 2'-FL |
| | $\times 2$ | S2B | - |
| Growth & death probability | $\times 0.1$ | S2C | Many others species are also abundant |
| | $\times 10$ | S2D | - |
| Probability of random new populations | $\times 0$ | S2E | - |
| | $\times 10$ | S2F | - |
| Diffusion | $\times 0.2$ | S2G | - |
| | $\times 5$ | S2H | - |
| Initial oxygen | $\times 0$ | S2I | Butyrate producers are also abundant without 2'-FL |
| | $\times 10$ | S2J | - |
| Quantity of mucin released | $\times 0$ | S2K | - |
| | $\times 5$ | S2L | *Bifidobacterium* nearly extinct without 2'-FL |
| GOS instead of 2'-FL | N/A | S3 | Higher abundance of *B. bifidum* and *B. breve* ($p < 0.001$)<br>Lower abundance of butyrate producers ($p < 0.001$) |
| Mucin replaced with non-fucosylated mucin | N/A | S4 | Similar outcomes |
| ATP + acetate as proxy for biomass | N/A | S5A&E | Similar outcomes, but *Lactobacillus gasseri* could never grow |
| Acetate as proxy for biomass | N/A | S5B&F | Microbiota consists nearly entirely of *B. longum* |
| ATP + pyruvate as proxy for biomass | N/A | S5C&G | *B. vulgatus* extinct |
| ATP + ketoglutarate as proxy for biomass | N/A | S5D&H | *B. vulgatus* extinct |

We concluded that the model was robust for an alternative prebiotic oligosaccharide, a non-fucosylated mucin, changes to the public goods production rate, and several other parameter changes. However, the assumed biomass reaction greatly impacted the predictions of the model. We decided not to continue with the alternative biomass proxies. Proxies 2-4 led to unrealistic outcomes. Alternative biomass proxy 1 led to predictions that were no worse than those with the default biomass reaction, but also did not match the *in vivo* data better. We proceeded to use the biomass function that required only ATP, as the resulting microbiota composition adequately matched *in vivo* observations of microbiota composition while allowing for growth of all GEMs [17,32]. We will discuss potential future improvements to the biomass reaction in the discussion section ‘Current and future extensions’.

### 2.4. B. bifidum produces cross-feeding metabolites that may allow it to be exploited by other bacteria

As we observed a bimodal distribution of mucin consumption in the model simulations with 2’-FL and public goods 2’-FL metabolism (Fig. 2B), we hypothesized that both a different metabolism and a different bacterial composition could be associated with the high and low mucin consumption simulation results. We next investigated why the model with public goods metabolism of 2’-FL led to (1) a lower mucin consumption in some simulations, but not in others, (2) a high *B. longum* abundance, and (3) a low *B. bifidum* abundance.

To gain insight into the potential mechanisms driving low versus high mucin consumption, we visualized the metabolic interactions in the final 60 timesteps of a simulation without 2’-FL (Fig. 3A), of a simulation with public goods metabolism of 2’-FL and low mucin consumption (Fig. 3B) and of a simulation with public goods metabolism of 2’-FL and high mucin consumption (Fig. 3C).

**Figure 3.**
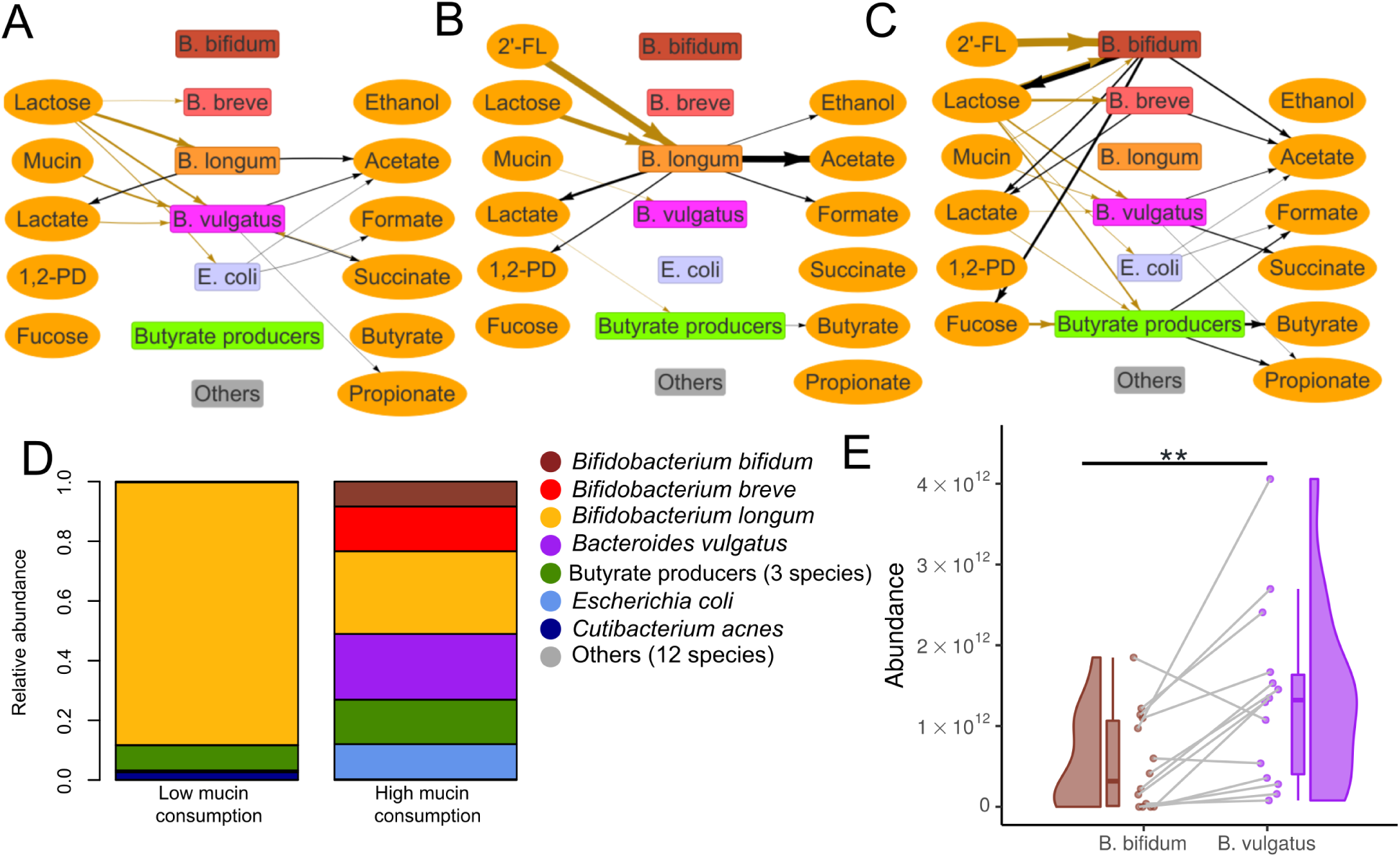
High consumption simulations show complex cross-feeding but higher abundance of *B. vulgatus* than *B. bifidum*. (A,B,C) Visualisation of metabolic interactions in timesteps 10020 to 10080 (last 3 hours) of a sample simulation (A) with 211 µmol lactose per 60 timesteps (B) with additional 211 µmol 2’-FL per 60 timesteps, public goods 2’-FL metabolism, and low mucin consumption (C) with additional 211 µmol 2’-FL per 60 timesteps, public goods metabolism for 2’-FL, and high mucin consumption. Line width is scaled with the flux per metabolite over the last 60 timesteps, multiplied by the carbon content of the molecule, with a minimum threshold of 100 µmol atomic carbon. (D) Average relative abundance of bacterial species in the condition with public goods 2’-FL metabolism, split by low (<50%) or high (>50%) mucin consumption at the end of 21 days. n=16&n=14, respectively. (E) Absolute abundance of *B. bifidum* and *B. vulgatus* in the simulations with public goods 2’-FL metabolism and high mucin consumption. Data from the same simulation is connected with a line. n=14. NS: Not significant, *: p<0.05, **:p<0.01, ***:p<0.001

The network visualisations revealed notable differences between these three situations. In the control simulation without 2’-FL a complex community formed, with high mucin consumption and extensive cross-feeding between *B. longum* and *B. vulgatus* (Fig. 3A). The simulation with 2’-FL and low mucin consumption showed less diversity: it became dominated by 2’-FL consuming *B. longum* and some non-mucin consumers, which were predicted to cross-feed on lactate (Fig. 3B). The high mucin consumption simulation with 2’-FL had a distinct complex community, in which *B. bifidum* consumed 2’-FL and many species cross-fed on the lactose produced as a public good by *B. bifidum* (Fig. 3C). *B. longum* was only present at a very low abundance in this simulation. From these networks we concluded that different communities arose in the simulations with high mucin consumption compared to the simulations with low mucin consumption.

To investigate the bacterial species associated with these communities in the model we cat-egorized the relative abundances of the simulations according to the rate of mucin consumption (Fig. 3D). Low mucin consuming simulations (<50% of mucin consumed, n=16) were consis-tently dominated by *B. longum*. High mucin consuming simulations (>50% of mucin digested, n=14) instead showed a higher abundance of *B. bifidum*, but also more *E. coli* and *B. vulgatus*. Interestingly, *B. bifidum* was typically not the dominant species in either set of simulations. We hypothesized that other species were taking advantage of the public good 2’-FL metabolism by *B. bifidum*, and proceeded to analyse the relationship between *B. bifidum* and the other species. In support of our hypothesis, *B. vulgatus* was more abundant than *B. bifidum* (paired samples Wilcoxon test, p=0.02) and also positively correlated (Spearman correlation,p<0.01,r=0.68) (Fig. 3E). This implies that *B. vulgatus* benefited from a higher abundance of *B. bifidum*. *B. vulgatus* was the only species within the high mucin consumption simulations to be both more abundant and positively correlated (Fig. S6).

Altogether, within the simulations *B. bifidum* produced public goods that were consumed by other species, whereas*B. bifidum* was consistently less abundant than *B. vulgatus*, leading us to hypothesize that *B. vulgatus* exploited *B. bifidum*’s metabolism of 2’-FL.

### 2.5. B. bifidum is exploited when digesting 2’-FL but B. longum is not

We continued to examine the possible interactions between *B. vulgatus* and *B. bifidum*. In particular, we hypothesized that *B. vulgatus* could exploit *B. bifidum* only in the public goods model of 2’-FL metabolism. This would mean that *B. vulgatus* could not exploit *B. bifidum* without 2’-FL, nor with 2’-FL modelled without public goods metabolism.

To further investigate possible interactions between *B. vulgatus* and *B. bifidum* we studied a simplified model with only *B. vulgatus* and *B. bifidum* and no other species. We then performed new simulations using the same four conditions as those shown in Fig. 2. In simulations with only *B. bifidum* and *B. vulgatus*, both species became abundant in absence of 2’-FL (Fig. 4A). In presence of 2’-FL, and with the non-public good variant of 2’-FL metabolism, *B. bifidum* became much more abundant than *B. vulgatus* (Fig. 4A). However, with public goods 2’-FL metabolism *B. vulgatus* became much more abundant than *B. bifidum* (Fig. 4A), despite *B. vulgatus* not consuming 2’-FL itself. The network visualization showed that without public goods metabolism of 2’-FL, *B. bifidum* did not produce lactose as a public good (Fig. 4B). By contrast, with public goods metabolism of 2’-FL *B. vulgatus* fed on the lactose produced by *B. bifidum* from 2’-FL (Fig. 4C). We concluded that the model predicted that the public goods metabolism of 2’-FL caused *B. vulgatus* to become relatively more abundant than *B. bifidum*, compared to the condition without 2’-FL.

**Figure 4.**
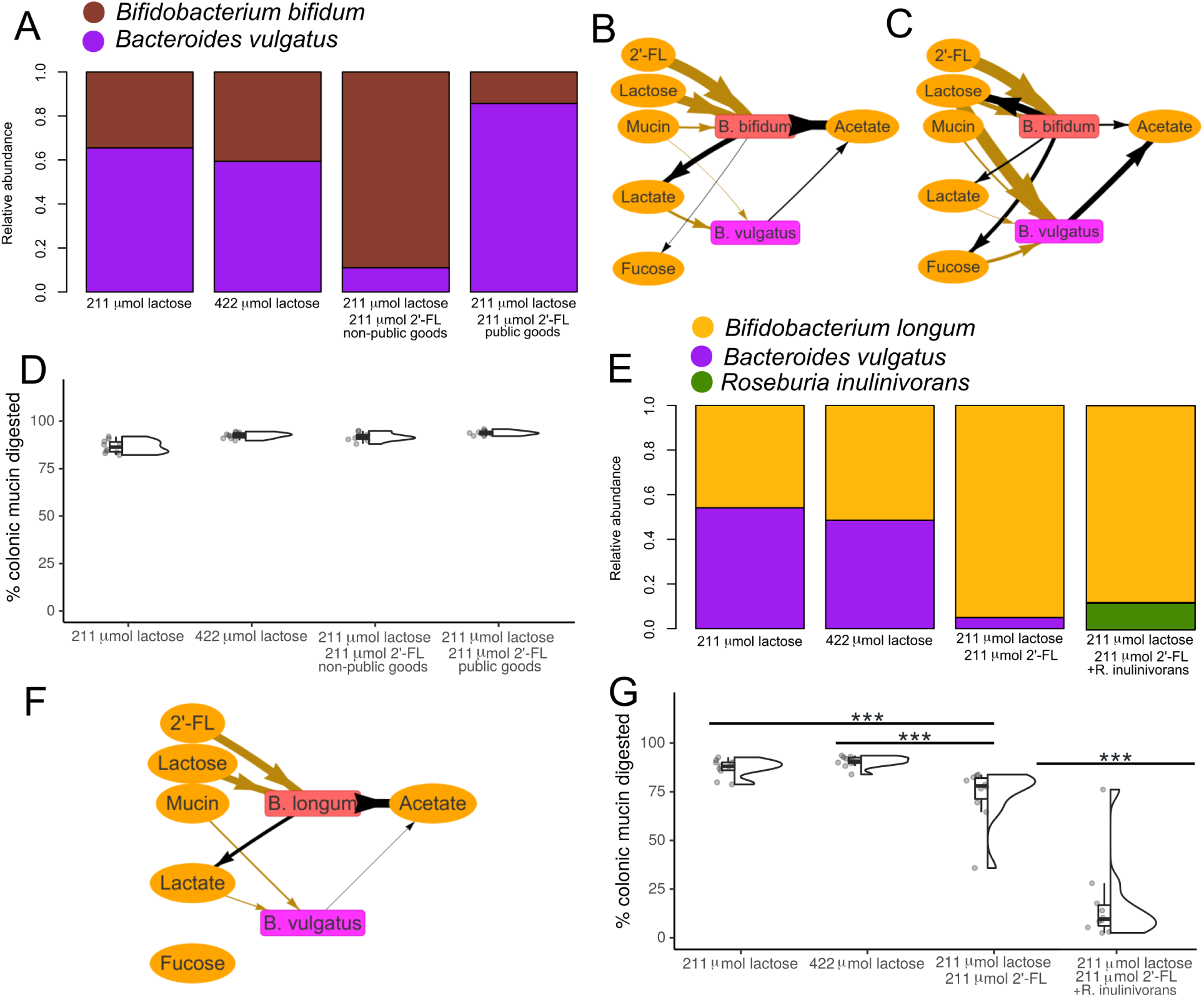
*B. bifidum*, but not *B. longum*, is exploited when digesting 2’-FL. (A) Average relative abundance of *B. bifidum* and *B. vulgatus*, with the following per 60 timesteps: 211 µmol lactose, 422 µmol lactose, 211 µmol lactose and 211 µmol 2’-FL with public goods 2’-FL metabolism, or 211 µmol lactose and 211 µmol 2’-FL without public goods 2’-FL metabolism, at the end of 21 days. n=10 per condition (B,C) Visualisation of metabolic interactions in sample simulations with only *B. bifidum* and *B. vulgatus* for (B) the non- public goods metabolism and (C) the public goods metabolism. Line width is scaled with the flux per metabolite over the last 60 timesteps, multiplied by the carbon content of the molecule, with a minimum threshold of 100 µmol atomic carbon. Data from the last 3 hours, step 10020 to 10080. (D) Amount of colonic mucin digested by the microbiota as a percentage of total mucin released into the gut over the final 3 hours of the model, per condition of A. n=10 for each. (E) Average relative abundance of *B. longum* and *B. vulgatus*, with 211 µmol lactose, 422 µmol lactose, or 211 µmol lactose and 211 µmol 2’-FL per 60 timesteps, at the end of 21 days. The simulations of the bar furthest to the right also include *R. inulinivorans*. (F) Visualisation of metabolic interactions in a sample simulation with only *B. longum* and *B. vulgatus*. Line width is scaled with the flux per metabolite over the last 60 timesteps, multiplied by the carbon content of the molecule, with a minimum threshold of 100 µmol atomic carbon. Data from the last 3 hours, step 10020 to 10080. (G) Amount of colonic mucin digested by the microbiota as a percentage of total mucin released into the gut over the final 3 hours of the model, per condition of E. n=10 for each. NS: Not significant, *: p<0.05, **:p<0.01, ***:p<0.001 (E) *R. inulinivorans* (Fig. S7A&B, p>0.05). With *B. bifidum*, *B. vulgatus* and *R. inulinivorans* there was also no reduction in mucin consumption compared to the same condition without *R. inulinivorans* (Fig. S7C&D, p>0.05). We therefore concluded that *B. longum* could indeed reduce mucin consumption in the model with 2’-FL, in contrast to *B. bifidum*.

Mucin consumption in the limited consortium of *B. bifidum* and *B. vulgatus* was high in all conditions, regardless of their relative abundances, as both species consumed mucin (Fig. 4D). To confirm whether *B. longum* reduced mucin consumption in competition with *B. vulgatus* we repeated the conditions with lactose and non-public goods digestion of 2’-FL (conditions 1-3), but with only *B. longum* and *B. vulgatus*. As neither species could perform public goods 2’-FL digestion we did not repeat the simulations with the public goods model of 2’-FL digestion. In absence of 2’-FL (conditions 1&2), *B. longum* and *B. vulgatus* were approximately equally abundant (Fig. 4E). In presence of 2’-FL *B. longum* became consistently more abundant than *B. vulgatus* (Fig. 4E). *B. longum* did not create lactose from 2’-FL like *B. bifidum* did (Fig. 4F). In contrast to *B. bifidum*, mucin consumption with *B. longum* and 2’-FL was lower compared to the simulations without 2’-FL (p<0.01,Fig. 4G).

In the limited consortium of *B. longum* and *B. vulgatus* the reduction in mucin consumption with 2’-FL was small, and *B. vulgatus* was still consistently present. We observed that *B. vulgatus* cross-fed on lactate produced by *B. longum* (Fig. 4F). We hypothesized that the consumption of lactate may have allowed *B. vulgatus* to survive in this limited consortium, but that this did not occur in the full consortium (Fig. 3D). We next added *Roseburia inulinivorans*, a lactate consumer from the full consortium, to see if it could outcompete *B. vulgatus* and so lower mucin consumption. *R. inulinivorans* indeed outcompeted *B. vulgatus* (Fig. 4E) and mucin consumption was indeed lower (Fig. 4G, p<0.01). To control for a possible effect of *R. inulinivorans* on mucin consumption without 2’-FL or without *B. longum* we performed several additional simulations. With *B. longum*, *B. vulgatus* and *R. inulinivorans* as species and without 2’-FL there was no reduction in mucin consumption compared to the same condition without

In conclusion, the model predicts that *B. bifidum* is exploited by *B. vulgatus* when it grows on 2’-FL. This happens because of the public good metabolism of *B. bifidum*. The model also predicts that when *B. longum* grows on 2’-FL it is not exploited by *B. vulgatus*, because *B. longum* does not produce public goods. This allows *B. longum* to outcompete *B. vulgatus*. Because *B. longum* does not consume mucin, mucin consumption is lower when it outcompetes the mucin consumer *B. vulgatus*. *B. longum* metabolites can also feed a non-mucin consuming lactate consumer that can outcompete *B. vulgatus*, further lowering mucin consumption.

## 3. Discussion

To provide a mechanistic explanation for why breastfeeding modifies the microbiota in such a way that the microbiota consumes less mucin we created a multiscale mathematical model in this study. Concretely, the model predicts that both the human milk oligosaccharide 2’-FL and galacto-oligosaccharides stimulate non-mucin consuming species at the expense of mucin consumers, thus suggesting a plausible mechanism for the potentially beneficial effect of human milk on mucin consumption by the microbiota [6]. The mechanism predicted by the model is as follows: oligosaccharides are consumed by *B. longum*, enabling it to outcompete mucin consuming species. *B. bifidum*, which consumes both mucin and oligosaccharides, will lose the competition against *B. longum* in a complex community, because of its extracellular metabolism of oligosaccharides. This extracellular metabolism produces intermediate digestion products which function as public goods, and allow competitors to consume them. *B. longum* digests oligosaccharides intracellularly and, therefore, does not produce public goods, and so is not sensitive to exploitation.

### 3.1. Comparisons with experimental data

Several *in vivo* and *in vitro* observations agree with our model predictions. Our model predicted that *B. bifidum* can consume mucin (Fig. 1C&D). In addition, when the model included public goods metabolism of mucin, *A. hallii* could cross-feed on the public goods that *B. bifidum* produced from mucin, in particular galactose (Fig. 1D). These predictions agree with experimental observations: *B. bifidum* has been shown to grow on mucin and *A. hallii* has been shown to grow on digestion products of *B. bifidum*, including galactose, *in vitro* [30]. When the model included public goods metabolism of 2’-FL, the model predicted that *B. bifidum* produces lactose and fucose from 2’-FL which can be taken up by other species (Fig. 3C&4C). This is in line with *in vitro* observations [7,41]. Furthermore, the model predicted that 2’-FL will stimulate the growth of *B. longum* in the infant gut microbiota (Fig. 2C). This is a consequence of *B. longum* consuming 2’-FL intracellularly, thereby not producing public goods that could be used by competitors (Fig. 2C & 4E). In agreement with this prediction, a high abundance of *B. longum* has also been observed *in vivo* as an effect of 2’-FL supplementation [18] or breastfeeding [31]. The model also predicted that the microbiota consumes less mucin in the presence of 2’-FL (Fig. 2D). In the model this is a result of the non-mucin consumer *B. longum* getting a competitive advantage over mucin consumers by 2’-FL (Fig. 2C). In agreement with this, less mucin is digested in *in vitro* fermentations using fecal samples of breastfed infants compared to fermentations using stool samples of formula-fed infants [6]. Within the simulations with 2’-FL, the model also predicted that a high abundance of *B. longum* reduces mucin consumption by the microbiota in the presence of 2’-FL (Fig. 3D). In agreement with this prediction, high abundance of *B. longum* is correlated with reduced mucin consumption *in vivo* in breastfed infants [15].

The simulations also predicted that in the presence of 2’-FL the public goods produced by *B. bifidum* are consumed by *B. vulgatus*, which prevents *B. bifidum* from becoming abundant. In agreement with this prediction, *B. bifidum* in the mouse gut has been shown to stimulate a, as of yet undetermined, species within the Bacteroidaceae, the family that contains *B. vulgatus* [42]. No *in vitro* research is available on the competition or cooperation between *B. bifidum* and *B.vulgatus* in isolation. However, there are some *in vitro* studies on competition between *B. bifidum* and the non-HMO consumer *Bifidobacterium breve* when fed with HMOs [16,21]. These studies reported contradictory results: *B. breve* became less abundant than *B. bifidum* in a short-term *in vitro* experiment [16], whereas it became more abundant than *B. bifidum* in a chemostat experiment with a continuous input flow of 2’-FL and output flow of metabolites [21]. The flow rate seems to explain the latter results. In the chemostat experiment containing a co-culture of *B. bifidum* and *B. breve*, the two species together were much less abundant after the flow rate was reduced [21]. However, in a culture of *B. bifidum* alone, a reduced flow rate only had a minor effect on the bacterial abundance [21]. This suggests that the flow rate particularly impacts bacterial interactions. The authors hypothesised that reduced substrate availability was the main cause of the observed reduction in bacterial abundance, but this does not explain why monocultures were less affected than co-cultures [21]. Interestingly, *in vitro* research on public goods production and consumption by two *Bacteroides* species actually found that high flow rates decreased public goods consumption and total abundance [43]. As both high and low flow rates caused a low total abundance and public goods consumption, intermediate flow rates may be important for bacterial abundance and public goods consumption in these systems. Computational modelling of a wide range of 2’-FL and public goods concentrations may help determine what factors are important in the outcome of bacterial competition in the infant gut.

### 3.2. Public good interactions in other microbial systems

The model predicts a crucial role for public goods metabolism and cross-feeding on public goods in determining the abundances of major species. Public goods mechanics have been previously studied *in vitro* in budding yeast (*S. cerevisiae*)[23]. Some yeast cells break down sucrose into glucose and fructose extracellularly with the enzyme invertase, but lose 99% of the glucose produced to diffusion, which then functions as a public good [23]. This leads to a dynamic community of invertase producers and non-producers [23,44]. The producers generate enough public goods to feed both themselves and a non-producing population. Co-existence is maintained because of a ‘snowdrift game’ where the less common strategy (producing or not producing) has an advantage over the more common strategy [23]. In the yeast system, the ratio of producers to non-producers depends on the cost associated with production [23]. High cost or low efficiency of production leads to a smaller fraction of producers [23]. In our model, *B. bifidum* is the only producer of public goods from 2’-FL, and all other species are non-producers. Co-existence between *B. bifidum* and other species also occurred in most of our simulations, implying that our model is somewhat similar to the system studied in yeast [23]. However, the assumptions and predictions of our model also differ in several ways from the yeast system. In the first place, there is no direct cost associated with public good production for *B. bifidum* in our model, in contrast to the yeast system [23]. In yeast, the cells expend energy to create invertase, which is modelled by directly decreasing their growth [23]. The yeast system predicts that producers are more abundant than non-producers when public goods production has low or no cost [23]. Despite the lack of a direct cost for the producer *B. bifidum* in our model, it was still less abundant than most other species. This may be because *B. bifidum* suffers an implicit cost from its metabolism, which gives it a competitive disadvantage: in contrast to *B. longum*, *B. bifidum* cannot actively transport 2’-FL into the cell [11]. Furthermore, *B. bifidum* is not able to consume fucose like *B. vulgatus* can [45]. An additional explicit metabolic cost for *B. bifidum*, as the producer has in the yeast model [23], may further reduce its predicted abundance. However, this cannot change the model prediction that *B. bifidum* is much less abundant than *B. longum*. Secondly, in contrast to the yeast system where digestion of sucrose is always extracellular, *B. longum* digests 2’-FL intracellularly. A similar ecological role has been coined the ‘loner’ type in *in vitro* cultures of *Pseudomonas aeruginosa* strains [46], where loners consume the substrate by themselves without producing public goods. In the *P. aeruginosa* system, loners exist next to producer and non-producer types. The loner type outcompetes non-producers, because the loners can obtain more substrate. The producers outcompete the loners because they can obtain even more substrate, as extracellular metabolism is more efficient [46]. ‘Loners’ have also been created through genetic modification in an *in vitro* budding yeast community [47]. As expected, yeast loners outcompete non-producers, but in contrast to *P. aeroginosa*, they also outcompete producers. In addition, yeast loners died out after they outcompeted producers. Our model predicts that, like the loner type in yeast, *B. longum* typically outcompetes both non-producers and producers (Fig. 2C). However, in our model, *B. longum* does not die out, as its metabolism is efficient enough to maintain a large population in the absence of a producer (Fig. 3B&D). Thus, neither the yeast nor the *P. aeruginosa* ‘loner’ types are an exact equivalent to the role the model predicts for *B. longum*. A third difference between the yeast system and the model predictions is that in the yeast system producers are more abundant when public good production is more efficient, due to a higher sucrose concentration. In contrast, the outcome of our model did not depend strongly on the amount of public goods produced per timestep by each population (Fig. S1). The lack of dependence on the rate of public goods production may be because of several reasons. The presence of other food sources, such as lactose, may dampen the effect of variation in public good availability in our model. Alternatively, *B. bifidum* may be so limited by its metabolism that more efficient public goods production cannot compensate sufficiently to significantly increase its abundance. Finally, diffusion may cause the local concentration of 2’-FL to become too low to allow for more public goods to be produced. Further study could determine what factor is decisive in our system.

### 3.3. Comparison to the infant gut microbiota composition

We can compare the model predictions for the relative abundances of species in the infant gut with available *in vivo* data. Our model predicts which species from our selection of 21 species become abundant in the infant gut in response to specific interventions. The simulations with lactose, as the only carbon source, i.e. without 2’-FL, predicted that *Escherichia*, *Bacteroides* and *Bifidobacterium* become the most abundant genera, in agreement with *in vivo* data on infants at the age of three weeks [17,32]. The model simulations with either GOS or 2’-FL also predicted a high abundance of *Bifidobacterium*, which matches *in vivo* data [18,31,48]. However, various discrepancies exist between the model predictions and *in vivo* data. Firstly, the model predicts a gut microbiota composition typically consisting of only a few species (Fig. 2C), whereas *in vivo* measurements have shown that the gut microbiota consists of dozens of species [32]. Most of these species have a low abundance *in vivo*, but they may still influence the dynamics of the system as a whole [17,49]. The inclusion of more species, a more extensive portrayal of bacterial metabolism combined with more complex nutritional input into the model will most likely further increase the accuracy and representativeness of the model. Secondly, the model does not predict a dominance of Bacillota, such as *Lactobacillus* or *Streptococcus* species, in any of the simulations (Fig. 2C), in contrast with *in vivo* data where such dominance is sometimes seen (e.g. in 18% of infants [50]). Although it is unclear why this is seen *in vivo*, there is a positive correlation with a lower gestational age and a defective mucin barrier [51,52]. The lower gestational age in particular may lead to a different initial composition, such as a composition without *Bifidobacterium*. These are factors that we did not include in the model. Lastly, the model does not reproduce differences in *Bifidobacterium* abundance as a result of mucin fucosylation (Fig. S4), as has been observed in an *in vivo* study [38]. It is unclear why different mucin fucosylation leads to different *Bifidobacterium* abundances [38]. Thus, it is also unclear what additions could be made to the model to improve these predictions.

### 3.4. Current and future extensions

A potential problem with the dynamic FBA modelling approaches that we use is their difficulty in reproducing *in vitro* results on relative growth rates of different combinations of bacteria [53,54]. Inaccurate predictions have been attributed to a combination of factors, primarily insufficient curation of reactions in GEMs, incorrect biomass reactions, and a lack of condition-specific constraints [53–56]. We have addressed these issues as follows. To address the insufficient curation of many GEMS, we created an additional curation pipeline that included checks for mass-balance and the absence of ATP production without substrate. We also performed extensive comparison with previous *in vitro* observation and corrected the presence or absence of important carbon uptake reactions (Methods section ‘Changes to genome-scale metabolic models’). As part of the model we imposed an enzymatic constraint on the FBA solutions (Methods section ‘Flux balance analysis approach’), which improves metabolic predictions by correctly predicting overflow metabolism in many situations where classic FBA would not [57]. We paid particular attention to the curation of the crucial *Bifidobacterium* GEMs, and ensured that the simulations correctly reproduced the substrate concentration dependent bifid shunt pathway [19,28]. To address the uncertain and arbitrary nature of biomass reactions included in the GEMs, our model uses a biomass reaction that only requires ATP, which simplifies the system and allows us to focus on curating the carbon metabolism of the bacteria. However, it does limit the scope of the predictions in our model, as we cannot incorporate interactions through, e.g. differences in amino acid availability, which may be important [58,59]. Finally, we addressed condition-specific constraints. We selected the simulated nutrient inputs in the model to be relevant to the infant gut situation, and constructed the ‘public goods’ system to better simulate extracellular metabolism. We have shown that the ‘public goods’ system allows the model to reproduce *in vitro* results that it otherwise would not capture (Fig. 1). However, as we discussed previously, there are a number of discrepancies between our model predictions and existing *in vitro* and *in vivo* data. We want to emphasize that it remains uncertain whether our model predictions are correct, and that *in vitro* and *in vivo* validation are essential.

The discrepancies between the model outcomes and existing *in vitro* and *in vivo* data are likely the result of the model’s incomplete and simplified representation of many aspects of the infant gut and the infant gut microbiota. We will now discuss several aspects in which the model is incomplete, and may be expanded in the future. At the species level, the selection used in the model is incomplete, as we do not include the vast majority of bacterial species present in the early developing infant gut [17]. We also do not include any viruses, fungi or archaea, which may play a significant role in metabolic interactions in the infant gut [60]. Future versions of the model may include a larger consortium of species. On a metabolic level, FBA requires many assumptions regarding bacterial metabolism, growth rate and biomass production [61,62]. To get insight into how the model predictions depended on our metabolic assumptions we performed further parameters variation after generating our main results. We examined the effect of different biomass reactions on the model predictions (Fig. S5) and found that the predictions were similar when the biomass reaction included acetate in addition to ATP, but not when other biomass reactions were used. This shows that the model can reproduce many results when carbon is taken up and used for growth, but that it is sensitive to the specific manner in which carbon is used for growth. These differences are most probably caused by the differences in FBA solutions. These, in turn, depend on the GEMs. Further study of the internal reactions selected in the different FBA solutions may shed more light on why the system is so sensitive to these variations. Furthermore, only biomass reactions consisting of a single carbohydrate or a single carbohydrate plus ATP were tested. In reality, however, bacterial metabolism requires many more components, including amino acids, nucleotides and cell wall components. Ideally, all these should be included in the biomass reaction. However, many of these these cellular components include nitrogen, which requires a good modelling of nitrogen availability and metabolism in the gut to be implemented in the system. The addition of such a system to future versions of the model will likely lead to more complex dynamics, as amino acids are also a cross-feeding substrate for some species [63] and the type and concentration of protein in infant formula influences microbiota composition [58,59]. Despite these discrepancies, the assumption that the ATP production rate corresponds with biomass production allowed the model to create accurate predictions for both the overall composition of the microbiota and the relative abundances of *B. longum* and *B. bifidum*.

The model is currently limited to a simplified representation of mucin. Only two different mucin structures are used in the model, but many more mucin structures exist. *In vivo* mucin structure is highly diverse and contains a greater number and higher complexity of mucins compared to the mucin structures used in our model [4,64]. A more realistic representation of mucin in the model would possibly result in more accurate model predictions about the developing gut microbiota when implemented. Furthermore, although human milk contains about 200 HMO structures, we focused our modeling on 2’-FL, as 2’-FL is the most abundant oligosaccharide in most human milk [65]. We also modelled GOS, as it is a common component of infant formulas [66]. If other HMOs were to be included in the model, this would lead to more complex metabolic interactions as these HMOs are digested using different enzymes [11]. A final limitation of the current model is that it does not include factors like gut pH or the influence of the host immune system [67,68]. These factors could also influence interactions and competition between bacteria in the infant gut, and thus influence the predictions of the model.

### 3.5. Potential health implications

The model predictions may have implications for the health and well-being of infants. Although it is currently unclear how the consumption of mucin by the gut microbiota directly impacts infant health, the observation that there is lower consumption of mucin in breastfed infants may suggest that mucin is important for normal growth and development of infants. This could be illustrated by the fact that breastfed infants are more protected against IBD and Crohn’s disease later in life than formula-fed peers [69], as both disorders have been associated with damage to the gastrointestinal mucin layer [70]. Strategies to prevent mucin loss, like nutritional interventions with 2’-FL or GOS as described in this paper, may be important in this light. Of course, other effects of breastfeeding, such as immune system modulation [71,72], may also play a role.

### 3.6. Outlook

Future versions of the model may consider a greater number of bacterial species, as well as a more complex representation of metabolism and metabolite sharing that could for instance include amino acids, nucleotides, and other oligosaccharides. The model can also be extended to simulate the adult gut. As dietary fiber becomes more abundant in the nutrition of infants after weaning [73], extracellular metabolism and public good dynamics may play an even larger role during childhood, adolescence and adulthood. Such extensions may provide further insights into the public good dynamics at work in the human gut.

## 4. Methods

### 4.1. Model overview

We have extended a multiscale spatial model of the infant gut. The model is based on our earlier models of the human microbiota [26] and of the infant microbiota in particular [28,29]. The major additions to this extension of the model are the inclusion of mucins and the public goods metabolism of mucins, GOS, and 2’-FL. In short, the model consists of a regular square lattice of 225 *×* 8 lattice sites (Fig. 2A). Each lattice site can contain a single population of a single species (Section 4.2, ‘Species composition’). Each species is represented by a genome-scale metabolic model, to which we have made some improvements to (Section 4.3, ‘Changes to genome-scale metabolic models’). Each lattice site can contain any number of nutrients and metabolites in any concentration. Metabolism determines growth (Fig. 2A-1) and is calculated for each population using flux balance analysis (FBA, section 4.4, ‘Flux balance analysis approach’). We have developed both a non-spatial (section 4.5, ‘Non-spatial model’) and a spatial version of the model (section 4.6, ‘Spatial model’). In the non-spatial model all populations, nutrients, and metabolites diffuse throughout the whole system every timestep. In the spatial model the populations, nutrients, and metabolites diffuse locally (Fig. 2A-2&3). To mimic movement through the gut all nutrients and metabolites also advect by one lattice site each timestep in the spatial model (Fig. 2A-4). Nutrients are input into the system at regular intervals to allow for metabolism (Section 4.7, ‘Nutrient input’). To represent the public-goods producing digestion of extracellular oligosaccharides, such as mucin, GOS, and 2’-FL, we have also created an additional model variant (Section 4.8, ‘Extracellular metabolism’). In this variant FBA is not used to calculate the breakdown of extracellular oligosaccharides. Finally, the model is initialised with randomly placed populations of all species, and the populations are allowed to grow and divide (Section 4.9, ‘Population dynamics’). This ultimately leads to a model that predicts a dynamic ecosystem with complex bacterial compositions and interactions, as we describe in the results.

### 4.2. Species composition

We selected the list of species to represent in the model (Table 1) from [17], using sheet 2 of their Table S3. We selected the 20 entries with the highest prevalence in vaginally delivered newborns. After removing two duplicate entries we selected a genome-scale metabolic model (GEM) of a species from each genus from the Virtual Metabolic Human database, generated in the AGORA project [74]. We then added an additional *Bifidobacterium breve* and *Bifidobacterium bifidum* GEM to represent the diversity of *Bifidobacterium* species in the infant gut [75]. We also added a GEM of *Roseburia inulinivorans*, as in our previous model [29]. *Roseburia* spp. have been shown to be prevalent butyrate producing bacteria in infants in other studies [76]. The populations of *Roseburia inulinivorans*, *Eubacterium hallii*, and *Clostridium butyricum* are pooled and listed as ‘Butyrate producers’ in the visualisations.

### 4.3. Changes to genome-scale metabolic models

We applied various changes and additions to the GEMs, as in our previous version of the model [29]. A full list of changed and added reactions is in table S1. We will highlight the more notable changes here. We disabled anaerobic L-lactate uptake for the *Bifidobacterium* and *E. coli* GEMs [77,78] and added a lactose symporter to *Anaerobutyricum hallii* [30], all *Bifidobacterium* GEMs [79], *Roseburia inulinivorans* [80], *Haemophilus parainfluenzae* [81], and *Rothia mucilaginosa* [82]. We also added galactose metabolism to *R. inulinivorans* [83] and *R. mucilaginosa* [82]. We have added metabolism of GOS to the *B. longum*, *B. breve* and *B. bifidum* GEMs and 2’-FL metabolism to the *B. longum* and *B. bifidum* GEMs. *B. longum* imports 2’-FL using an ABC transporter [84] and digests it to lactose and fucose inside the cell. *B. bifidum* breaks down 2’-FL to lactose and fucose using an extracellular fucosidase [7]. *B. longum* and *B. breve* import DP3 fractions of GOS intracellularly, and digest them to monosaccharides using beta-galactosidases [85]. *B. longum* and *B. breve* break down fractions longer than DP3 extracellularly using glycoside hydrolases [35,36]. *B. bifidum* digests all fractions extracellularly to lactose and galactose using extracellular beta-galactosidases [34]. We also added a fucosidase reaction to *B. bifidum* to allow it to remove fucose groups from mucin in line with the available literature [86].

### 4.4. Flux balance analysis approach

We use a modified version of FBA with an enzymatic constraint [61,87], as in previous model versions [28]. First, each GEM is converted to a stoichiometric matrix S. Reversible reactions are converted to two irreversible reactions, so that fluxes will be greater than or equal to 0. Reactions identified in the GEM as ‘exchange’, ‘sink’, or ‘demand’ take up or deposit metabolites into the environment. We assume that intracellular regulation occurs at a much faster rate than any extracellular dynamics, including population growth and spread and diffusion of extracellular metabolites. Subject to this separation of timescales, we can assume that all reactions are in internal steady state,

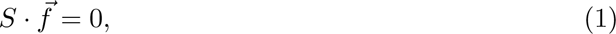

where 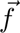 is a vector of the metabolic fluxes through each reaction in the network, in mol per time unit per population unit. Thus we apply a flux-balance analysis approach [61] to predict exchange fluxes as a function of extracellular concentrations.

Each exchange reaction that takes up metabolites from the environment *F_in_* is constrained by an upper bound *F_ub_* which represents the availability of metabolites from the environment. It is determined as follows,

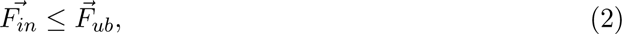

where 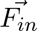 is a vector of fluxes between the environment and the bacterial population. 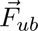 is a vector of upper bounds on these fluxes. 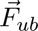 is set dynamically at each timestep t by the spatial environment at each lattice site *x⃗*:

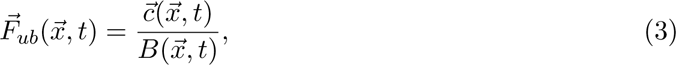

Where *c⃗* is a vector of all metabolite concentrations in mol per lattice site, *x⃗* is the location and *B*(*x⃗*, *t*) is the size of the local bacterial population [28]. 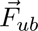 is set to 0 for any metabolite for which the public goods metabolism system is enabled for the local bacterial population (see section ‘Extracellular metabolism’).

The total flux through the network in each FBA solution is constrained by the enzymatic constraint *a*, in mol per time unit per population unit [28,87]. The enzymatic constraint represents the maximum, total amount of flux that can be performed per cell in each population:

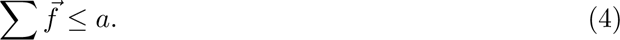

As both *f⃗* and *a* are given as a flux per population unit, this limit scales linearly with population size. Given these constraints, FBA uses linear programming to identify a solution that optimizes the objective function, ATP production. The solution consists of a set of input and output exchange fluxes 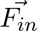(*x⃗*, *t*) and 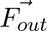(*x⃗*, *t*), and a growth rate *g*(*x⃗*, *t*). The exchange fluxes are taken as the derivatives of a set of partial-differential equations to model the exchange of metabolites with the environment. The size of the population increases proportionally to the growth rate in the FBA solution. Populations above 2 *·* 10^10^ bacteria do not perform metabolism to mimic quiescence at high densities.

### 4.5. Non-Spatial model

For the simulations of Fig. 1 we used a non-spatial version of the model to represent well-mixed *in vitro* conditions. All nutrients and metabolites are distributed uniformly across all lattice sites at the end of each timestep. All populations are moved to random locations at the end of each timestep. This eliminates spatial variation in metabolism. The change in concentration per lattice site is thus determined as follows,

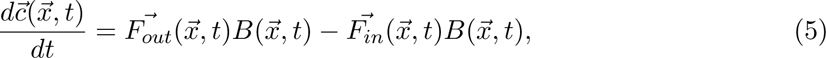

where 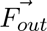(*x⃗*, *t*) is a vector of fluxes from the bacterial populations to the environment, in mol per time unit per population unit. All other parameters, including the lattice size, are as in the spatial model.

### 4.6. Spatial model

For all simulations after those of Fig. 1 we used a spatial version of the model. This version also used a regular square lattice of 225 x 8 lattice sites. In the spatial model mixing by colonic contraction is mimicked by diffusion. To each lattice site we apply the exchange fluxes as predicted by the FBA solution, yielding:

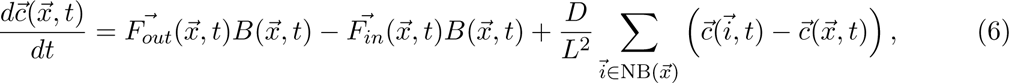

where *D* is the diffusion constant, *L* is the lattice side length, and *NB*(*x⃗*) are the four nearest neighbours.

Diffusion is applied to the metabolite concentrations on each lattice site at each timestep to represent mixing by colonic contractions. Metabolic diffusion is applied twice during each timestep. Each time it is applied, 14.25% of each metabolite diffuses from each lattice site to each of the four nearest neighbours. This causes a net diffusion each timestep of 6.3 *·* 10^5^ *cm*^2^/s. To mimic advection all metabolites except oxygen are moved distally by one lattice site every timestep. The transit time through the colon is approximately 11 hours in the model, corresponding with *in vivo* observations in newborn infants [88,89]. Metabolites at the most distal column of the lattice, the end of the colon, are removed from the system at each timestep. This represents the simulated feces.

### 4.7. Nutrient input

The secretion of mucin is mimicked as follows: each timestep a small concentration of mucin is added to the bottom-most row of the model. We used a core-2 mucin with one additional fucose, known in the VMH database as ‘MGlcn23 rl’, except for the simulation of Fig. S4. Here a core-2 mucin without additional fucose was used, known in the VMH database as ‘core2 rl’. In most simulations nutrients representing inflow from the small intestine are inserted into the first six columns of lattice sites every 60 timesteps, representing three hours, a realistic feeding interval for newborn infants [90]. Food intake contains 211 µmol of lactose (‘lcts’ in the VMH database) by default, a concentration in line with human milk [91], assuming 98% host uptake of carbohydrates before reaching the colon [25]. In some simulations 211 µmol of additional lactose, GOS, or 2’-FL is added. 2’-FL and GOS were not present in the VMH database. GOS are inserted as separate fractions of DP3, DP4, or DP5 based on analysis of the composition of Vivinal-GOS [33]. 64% is DP3, 28% is DP4 and 8% is DP5. Water (‘h2o’ in the VMH database) is provided in unlimited quantities.

### 4.8. Extracellular metabolism

We will now discuss the details of extracellular public-goods producing metabolism in our model. The GEMs of many species in our model contain reactions for the extracellular breakdown of oligosaccharides, such as mucins. We created a separate system for public goods-producing extracellular metabolism. In the simulations where this system was enabled, the oligosaccharides were excluded from FBA. Instead, 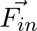 values for extracellular oligosaccharides were set directly by the environment as follows: The model applies each extracellular reaction at a rate of 2 µmol per 1*·*10^10^ population per timestep. This is entered into the sets of input exchange fluxes 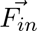 and output exchange fluxes 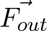. If two reactions can apply to a substrate, and insufficient substrate is available for each reaction to apply fully, each reaction is applied to half the remaining substrate. As no substrates have more than two reactions associated with them in our model, this ensures that breakdown is limited to the total amount of substrate available:

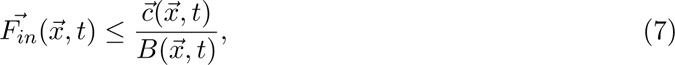

The change in concentration of metabolites is still determined by eq. 5 in the non-spatial model and eq. 6 in the spatial model.

Because the fluxes from the FBA and the separate extracellular metabolism are applied simultaneously they cannot interact within their own timestep. Diffusion and advection are applied before the products can be used by FBA. The public goods metabolism is applied to mucins in all simulations of the model, except those in Fig. 1C. The public goods metabolism is also applied to extracellular 2’-FL or GOS, and is noted as ‘public goods’ when used. All reactions related to 2’-FL and GOS breakdown are listed in S1 table. All mucin breakdown reactions are listed in table S2. In short, this approach lets us mimic the production of a larger variety and quantity of public goods from oligosaccharides than the Flux balance analysis approach does.

### 4.9. Population dynamics

The model is initialized by giving each lattice site a probability of 0.3 to generate a population of 5 *·* 10^7^ bacteria of a single random species. To this end, a GEM corresponding with this species (section ‘Species composition’) is associated with this lattice site. The exchange rates of metabolites for each population are calculated using FBA, based on the GEM, the enzymatic constraint *a*, its current population size *B*(*x⃗*, *t*) and the local concentrations of metabolites *c⃗*(*x⃗*, *t*). The outcome is applied to the environment (eq. 6) and the growth rate *g*(*x⃗*, *t*) to the local population size, as follows:

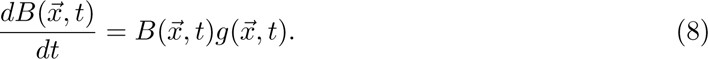

After initialisation, new bacterial populations can be created through reproduction (1) or introduction of new populations (2). To mimic reproduction (1), each population of at least 1 *·* 10^10^ bacteria (Table 3) creates a new population of the same species in an adjacent empty lattice site. Half the population size is transferred to the new population, so that biomass is conserved. To mimic introduction of populations (2) for each empty lattice site, with a probability of 0.00005, a species is selected at random, with equal probability for each species, and a population of this species is introduced into the lattice site. We initialize these populations at the same population size (*B*) as the initial populations in the model (Table 3). Finally, population are removed from the system with a probability of 0.0075 per population per timestep.

**Table 3.**
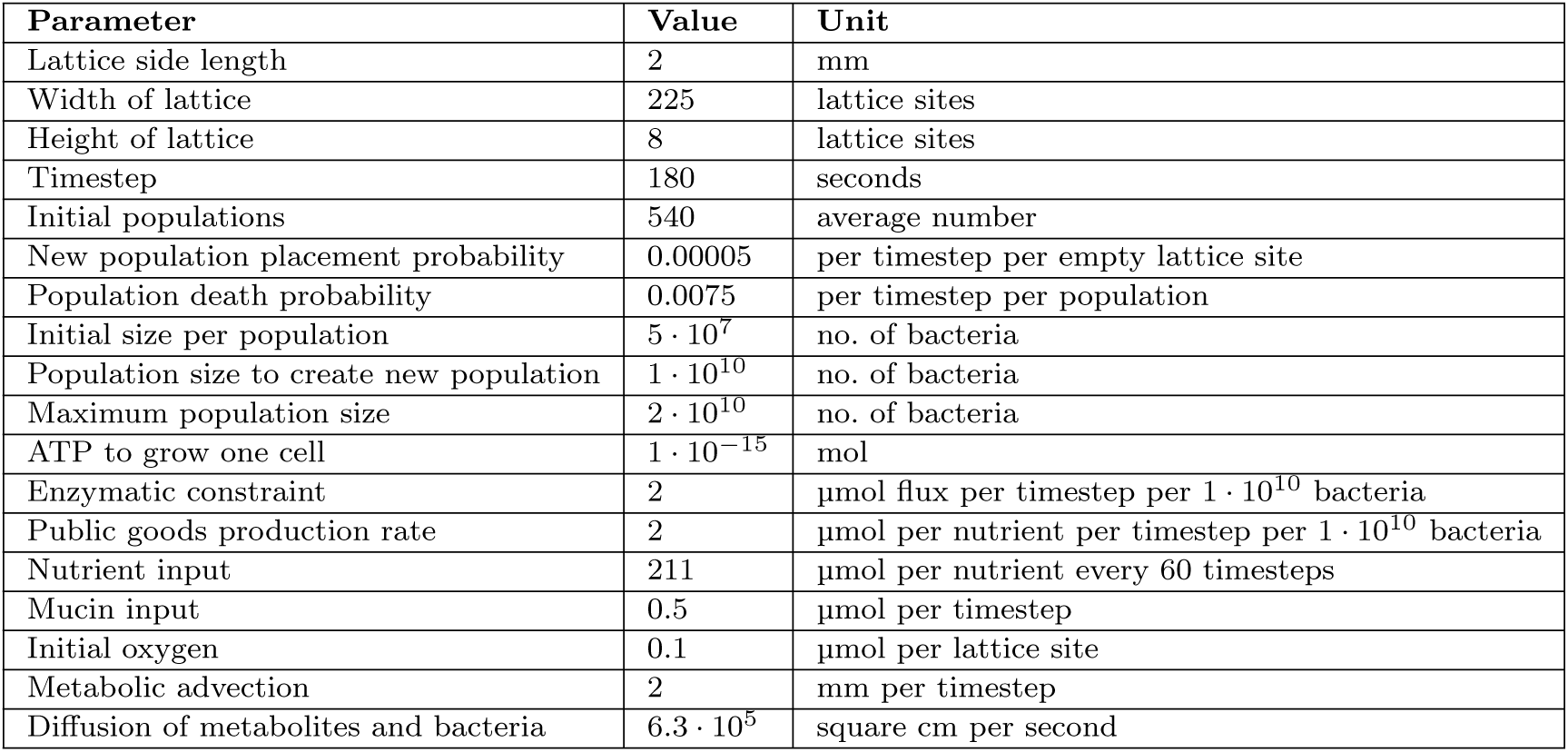
Parameters of the model.

| Parameter | Value | Unit |
| --- | --- | --- |
| Lattice side length | 2 | mm |
| Width of lattice | 225 | lattice sites |
| Height of lattice | 8 | lattice sites |
| Timestep | 180 | seconds |
| Initial populations | 540 | average number |
| New population placement probability | 0.00005 | per timestep per empty lattice site |
| Population death probability | 0.0075 | per timestep per population |
| Initial size per population | $5 \cdot 10^7$ | no. of bacteria |
| Population size to create new population | $1 \cdot 10^{10}$ | no. of bacteria |
| Maximum population size | $2 \cdot 10^{10}$ | no. of bacteria |
| ATP to grow one cell | $1 \cdot 10^{-15}$ | mol |
| Enzymatic constraint | 2 | $\mu\text{mol}$ flux per timestep per $1 \cdot 10^{10}$ bacteria |
| Public goods production rate | 2 | $\mu\text{mol}$ per nutrient per timestep per $1 \cdot 10^{10}$ bacteria |
| Nutrient input | 211 | $\mu\text{mol}$ per nutrient every 60 timesteps |
| Mucin input | 0.5 | $\mu\text{mol}$ per timestep |
| Initial oxygen | 0.1 | $\mu\text{mol}$ per lattice site |
| Metabolic advection | 2 | mm per timestep |
| Diffusion of metabolites and bacteria | $6.3 \cdot 10^5$ | square cm per second |

To mix the bacterial populations, the lattice sites swap population contents each timestep. We use a random walk algorithm based on Kawasaki dynamics [92], also used previously [26,28]. Each site is addressed in a random order, and the contents are swapped with a site randomly selected from the Moore neighbourhood. The contents consist of the bacterial population size *B*(*x⃗*, *t*) and the GEM. The swap only occurs if both the origin and destination site have not already swapped in this timestep. With this mixing method the diffusion constant of the bacterial populations is 6.3 *·* 10^5^*cm*^2^/*s*, equal to that of the metabolites. Bacterial populations at the most distal column, i.e. at the exit of the colon, are removed from the system. To increase the bacterial diffusion rate in the sensitivity analysis this process was executed five times, marking all sites as unswapped after each execution. To decrease the bacterial diffusion rate the number of swaps was limited to a fifth of the usual number of swaps.

### 4.10. Validity checks on FBA solutions

We performed a number of checks on the FBA solutions to ensure the model produces plausible predictions. We first checked whether the right species could grow on lactose. In line with *in vitro* literature all GEMs could grow on lactose, except *Veillonella disparans* [93], *Cutibacterium acnes* [94], *Eggerthella* sp. YY7918 [95], and *Gemella morbillorum* [96]. No GEM could grow without a substrate. We also checked each FBA solution for thermodynamic plausibility during the simulations using a database of Gibbs free energy values [97]. Values for 2’-FL, GOS, and mucin structures were generated using the values for their monosaccharides. All values assumed a pH of 7 and an ionic strength of 0.1 M, which are close to cytoplasmic values for common gut bacteria under acidic gut conditions [98–100]. Energy loss *l* in joules per timestep per population unit was recorded as follows, where *i* are metabolites, *F* is the exchange flux rate in mol per timestep per population unit and *E* contains the Gibbs free energy in joules per mol for each metabolite,

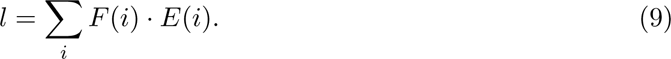

In the simulations of Fig 2 with 211 µmol of lactose per 60 timesteps (n=30) 99.6% of all FBA outcomes had a lower or equal amount of Gibbs free energy in the output compared to the input. The remaining 0.4% of FBA solutions was responsible for 0.07% of total bacterial growth. In the simulations with an additional 211 µmol of 2’-FL per 60 timesteps and public goods metabolism of 2’-FL (n=30) 99.3% of all FBA outcomes had a lower or equal amount of Gibbs free energy in the output compared to the input. The remaining 0.7% of FBA solutions was responsible for 0.04% of total bacterial growth.

### 4.11. Parameters

Parameters of the system are listed in table 3. We estimate that the infant colon has a volume of 90ml [101,102]. This leads to a rough estimate on the order of 10^12^ bacteria in the newborn infant colon given an abundance per ml of around 10^10^ [103]. Values for free parameters were estimated and evaluated in the sensitivity analysis(Fig. S1&Fig. S2).

### 4.12. Implementation details

The model is implemented in C++ 11 with libSBML 5.18.0 for C++ to load GEMs and the GNU Linear Programming Kit 4.65 (GLPK) to solve the FBA problems. Random numbers were generated with Knuth’s subtractive random number generator algorithm [104]. Diffusion of metabolites was implemented using the Forward Euler method. The model is based on our own earlier models of the gut microbiota [26,28,29]. GEMs are sourced from the May 2019 update of AGORA, the latest at time of writing, from the Virtual Metabolic Human Project website (vmh.life). We used Python 3.6 to extract thermodynamic data from the eQuilibrator API (December 2018 update) [97]. All p-values were calculated with R 4.2.2. Unless noted otherwise p-values were calculated using the Mann-Whitney test. Model screenshots were made using the libpng16 and pngwriter libraries. Other visualisations were performed with R 4.2.2. Raincloud visualisations used a modified version of the Raincloud plots library for R [105].

## 5. Supplemental material

***S1 Table***

**S1table.csv**

A table of changes made to the AGORA models as a .csv file.

***S2 Table***

**S2table.csv**

A table of mucin reactions used for public goods metabolism as a .csv file.

---

**S1 Figure.**
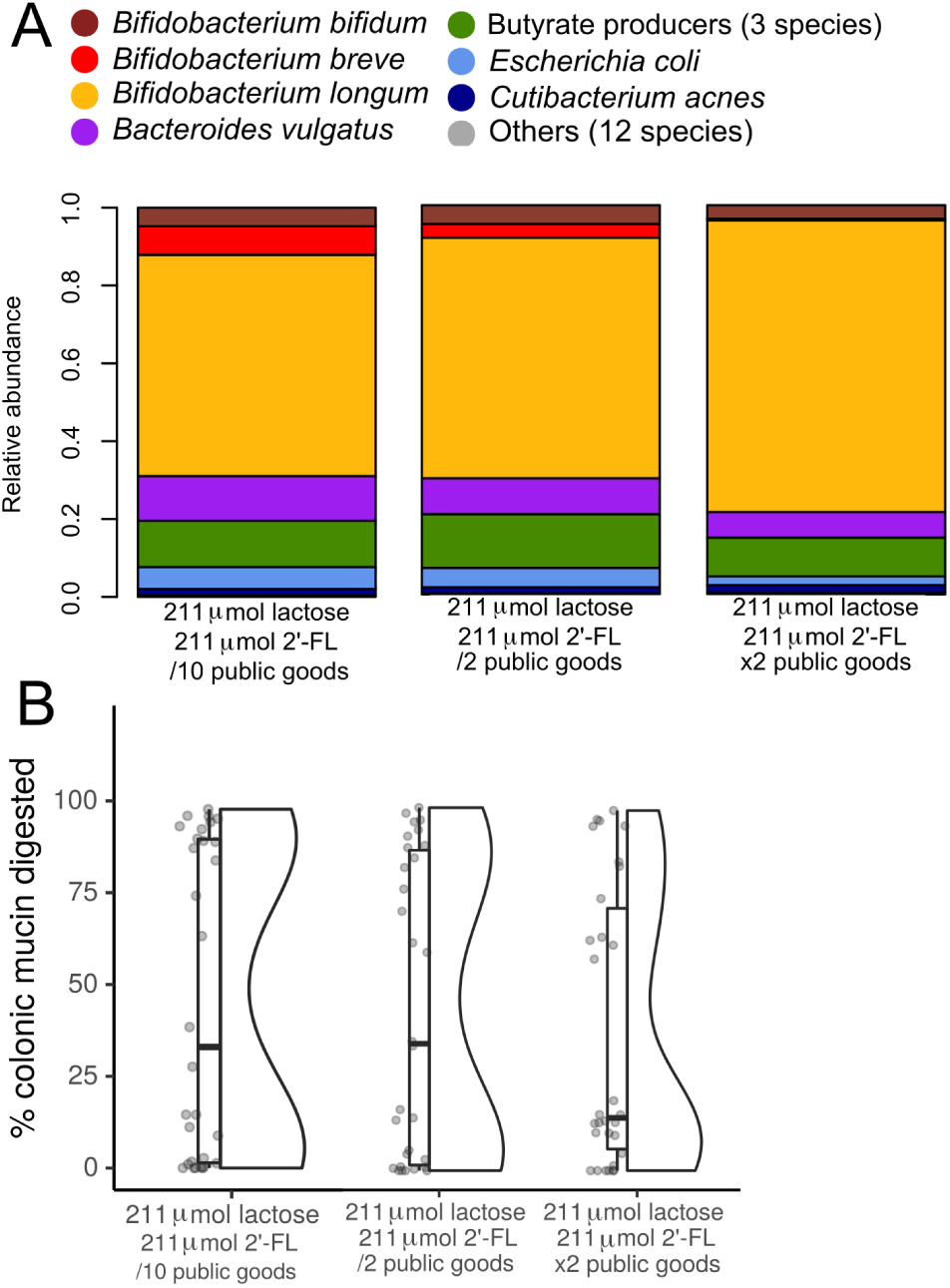
(A) Average relative abundance of bacterial species in the condition with 2’-FL with either the public good production rate reduced to a tenth, halved, or increased to double. n=30 for each condition. (B) Amount of colonic mucin digested by the microbiota as a percentage of total mucin released into the gut over the final 60 timesteps of the model, per condition corresponding to A. n=30 for each condition

**S2 Figure.**
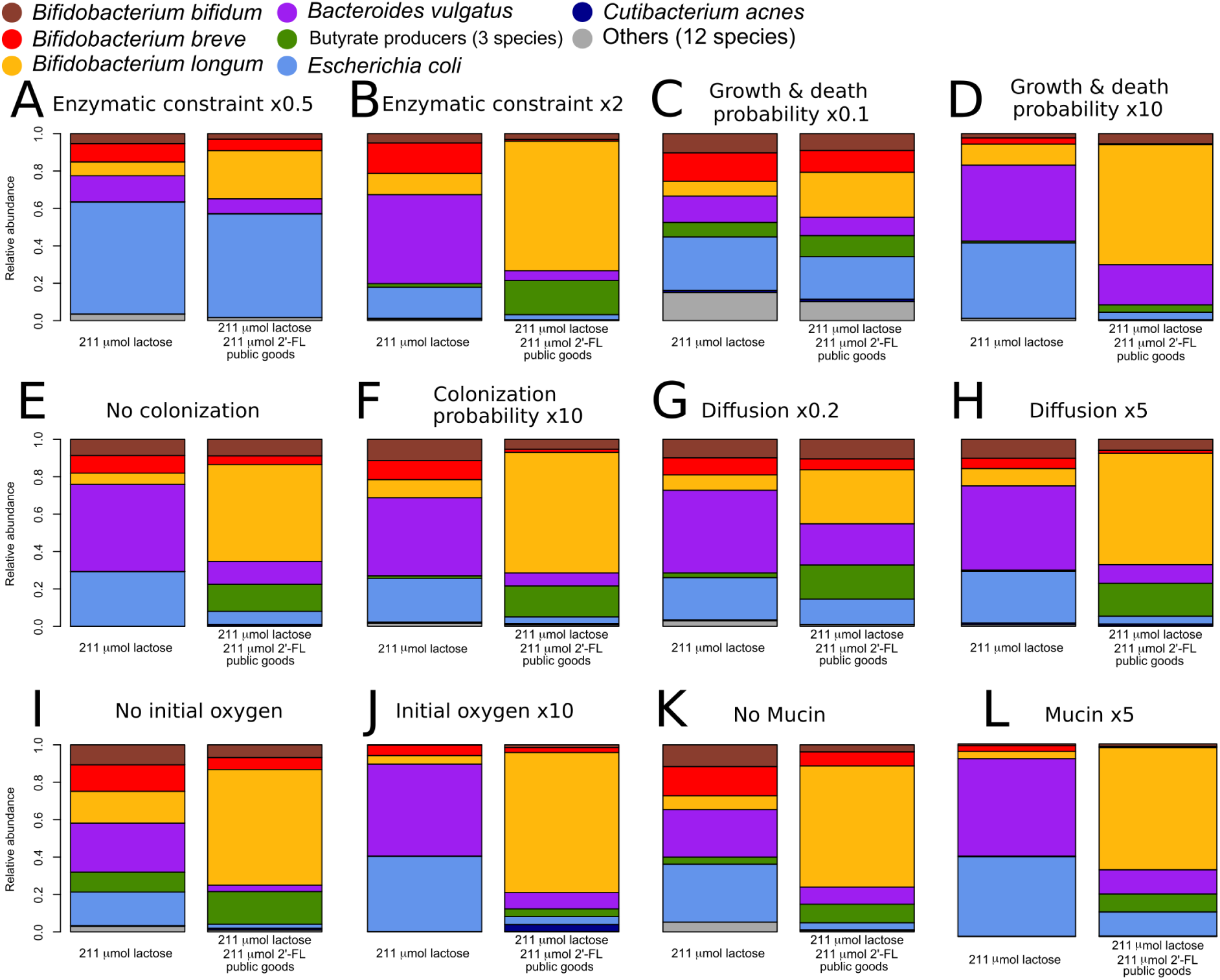
(A to J) Average relative abundance of bacterial species in the conditions with only lactose, or with 2’-FL and public goods 2’-FL metabolism, at the end of 21 days, with the following alteration from the baseline of Fig. 2A: (A) Enzymatic constraint loosened by a factor of 2, to 4 µmol flux per timestep per 1 *·* 10^10^ population (B) Enzymatic constrained tightened by a factor of 2, to 1 µmol flux per timestep per 1 *·* 10^10^ population (C) Growth decreased by a factor of 10, by increasing the ATP required to grow one bacteria to 1 *·* 10*^−^*^14^, with the death probability decreased to 0.00075 per population per timestep. (D) Growth increased by a factor of 10 by decreasing the ATP required to grow one bacteria to 1 *·* 10*^−^*^16^, with the death probability increased to 0.075 per population per timestep (E) Colonisation removed by setting the probability for new populations to be placed after initialization to 0 (F) Colonisation increased by a factor of 10 by setting the probability per empty lattice to acquire a new population to 0.0005 per timestep (G) Diffusion of both metabolites and bacteria decreased by a factor of 5 to 1.26 *·* 10*^−^*^6^ *cm*^2^/s (H) Diffusion of both metabolites and bacteria increased by a factor of 5 to 3.15 *·* 10*^−^*^5^ *cm*^2^/s (I) No initial presence of oxygen (J) Initial oxygen increased to 1 µmol per lattice site (K) No secretion of mucin (L) Secretion of mucin increased to 2.5 µmol per timestep. For each figure: n=30 for each condition, each simulation is weighed equally.

**S3 Figure.**
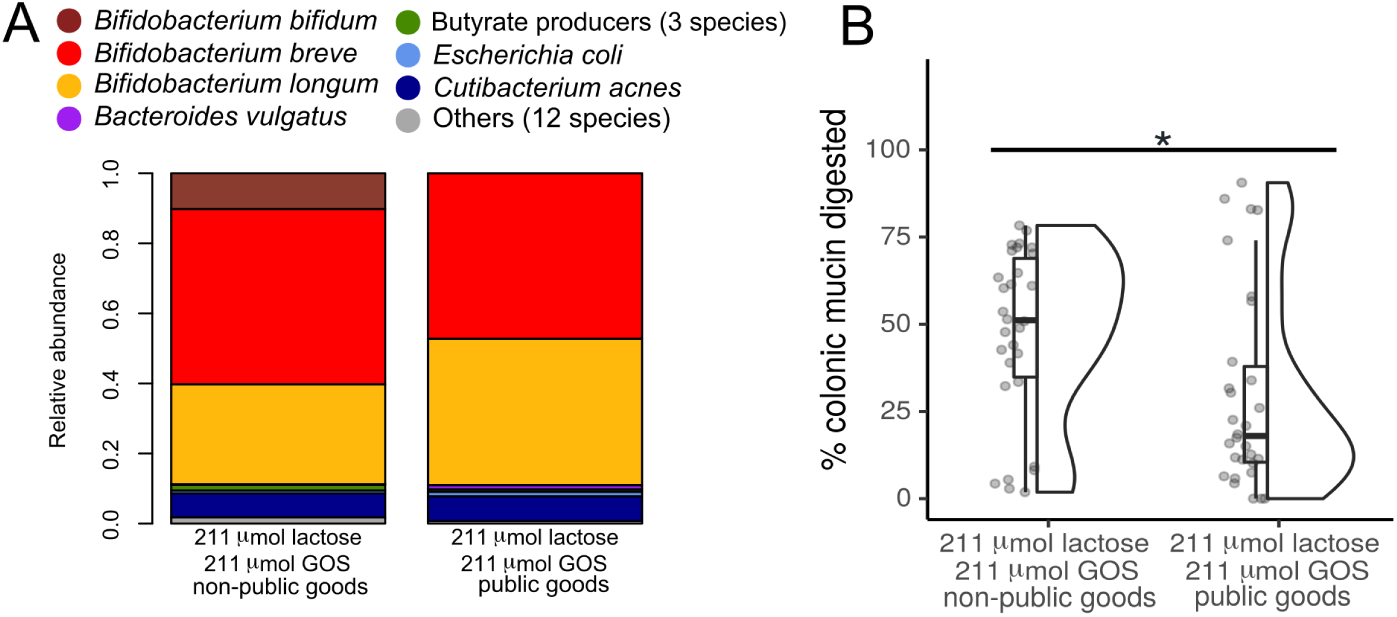
(A) Average relative abundance of bacterial species in the condition with GOS without public goods metabolism of GOS or GOS with public goods metabolism of GOS at the end of 21 days. n=30 for each condition. (B) Amount of colonic mucin digested by the microbiota as a percentage of total mucin released into the gut over the final 60 timesteps of the model, per condition. n=30 for each condition

**S4 Figure.**
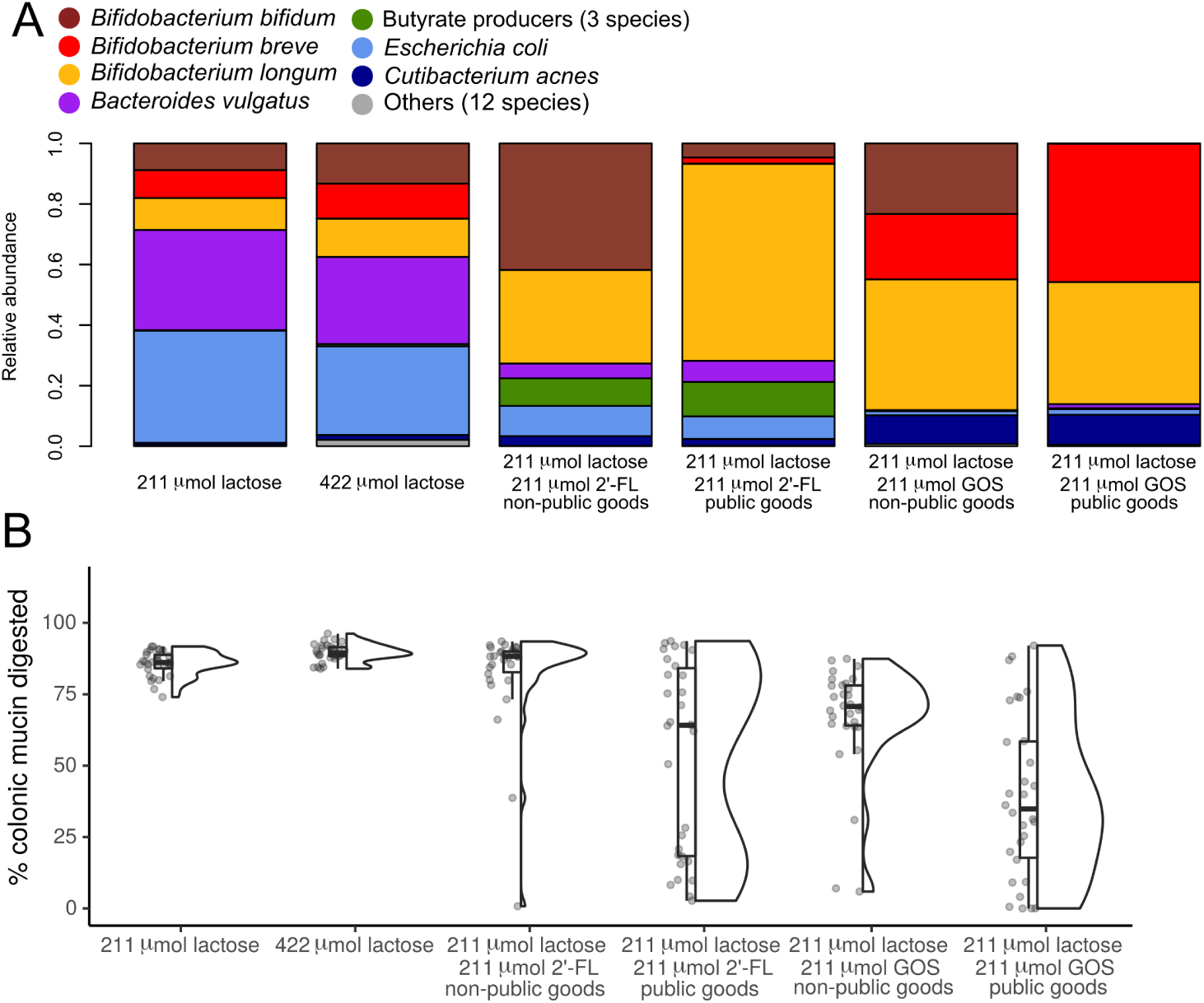
(A) Average relative abundance of bacterial species with non-fucosylated mucin instead of fucosylated mucin in each condition from Fig. 2 and Fig. S4 at the end of 21 days. n=30 for each condition. (B) Amount of colonic mucin digested by the microbiota as a percentage of total mucin released into the gut over the final 60 timesteps of the model, per condition. n=30 for each condition

**S5 Figure.**
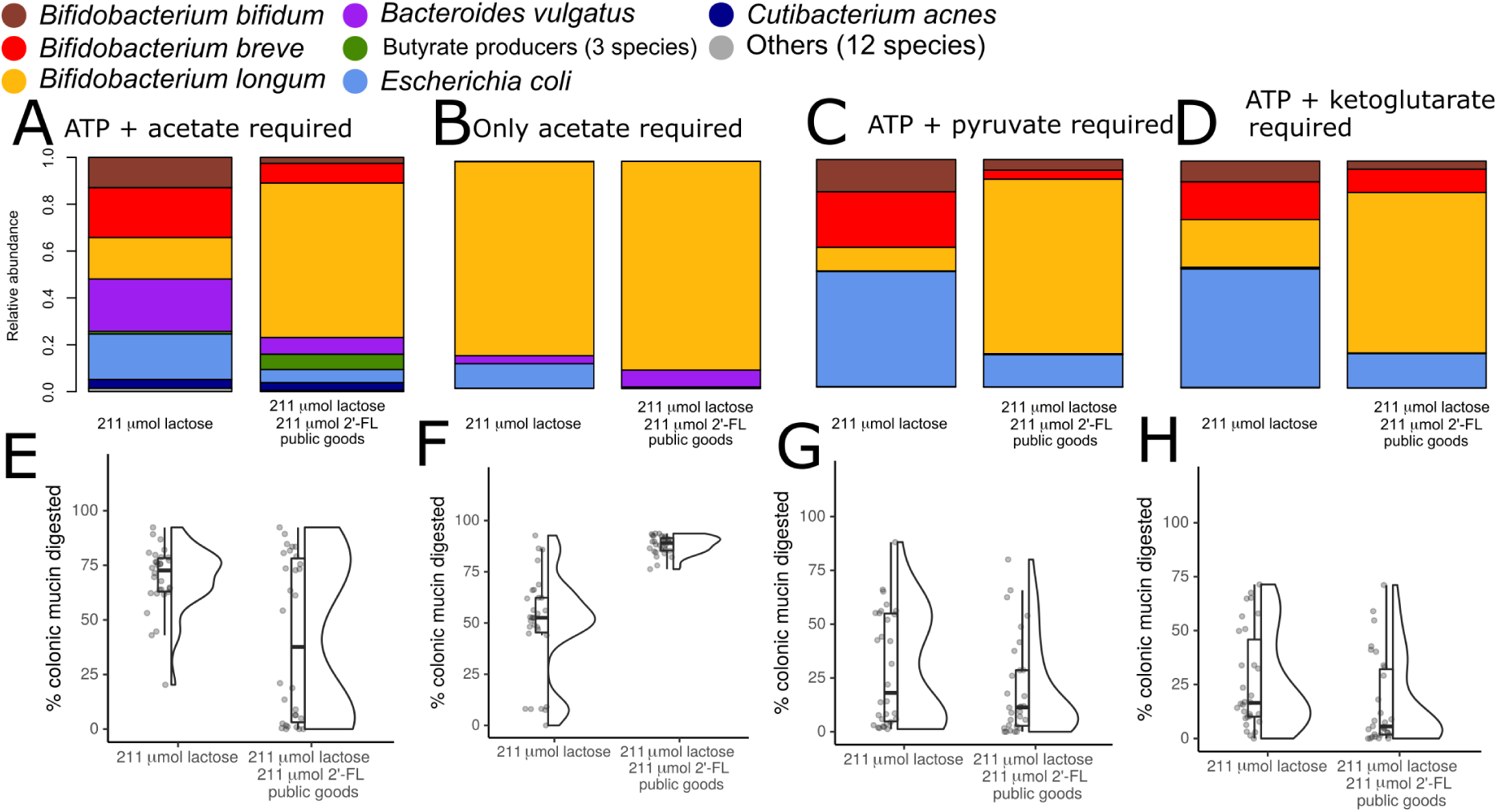
(A-D) Average relative abundance of bacterial species at the end of 21 days with the following alternative biomass reactions: (A)ATP + acetate (B) acetate (C) ATP + pyruvate (D) ATP + ketoglutarate n=30 for each condition. (E-H) Amount of colonic mucin digested by the microbiota as a percentage of total mucin released into the gut over the final 60 timesteps of the model, associated with A-D. n=30 for each condition

**S6 Figure.**
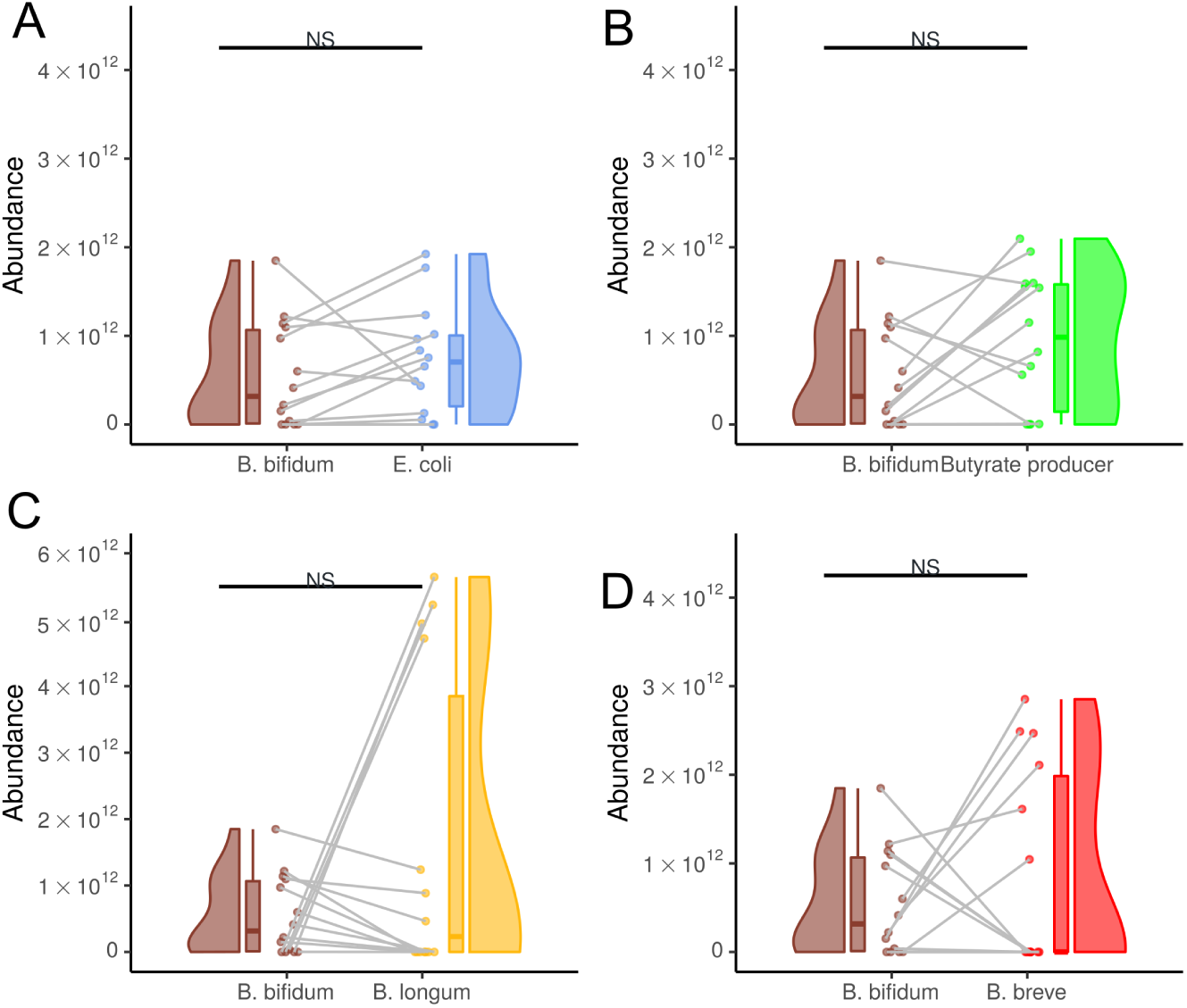
(A,B,C,D) Absolute abundance of *B. bifidum* compared to (A). *E. coli* (B) Butyrate producers (3 species) (C) *B. longum* (D) *B. breve* in the simulations with high mucin consumption of the public goods condition. Data from the same simulation is connected with a line. n=14. NS: Not significant

**S7 Figure.**
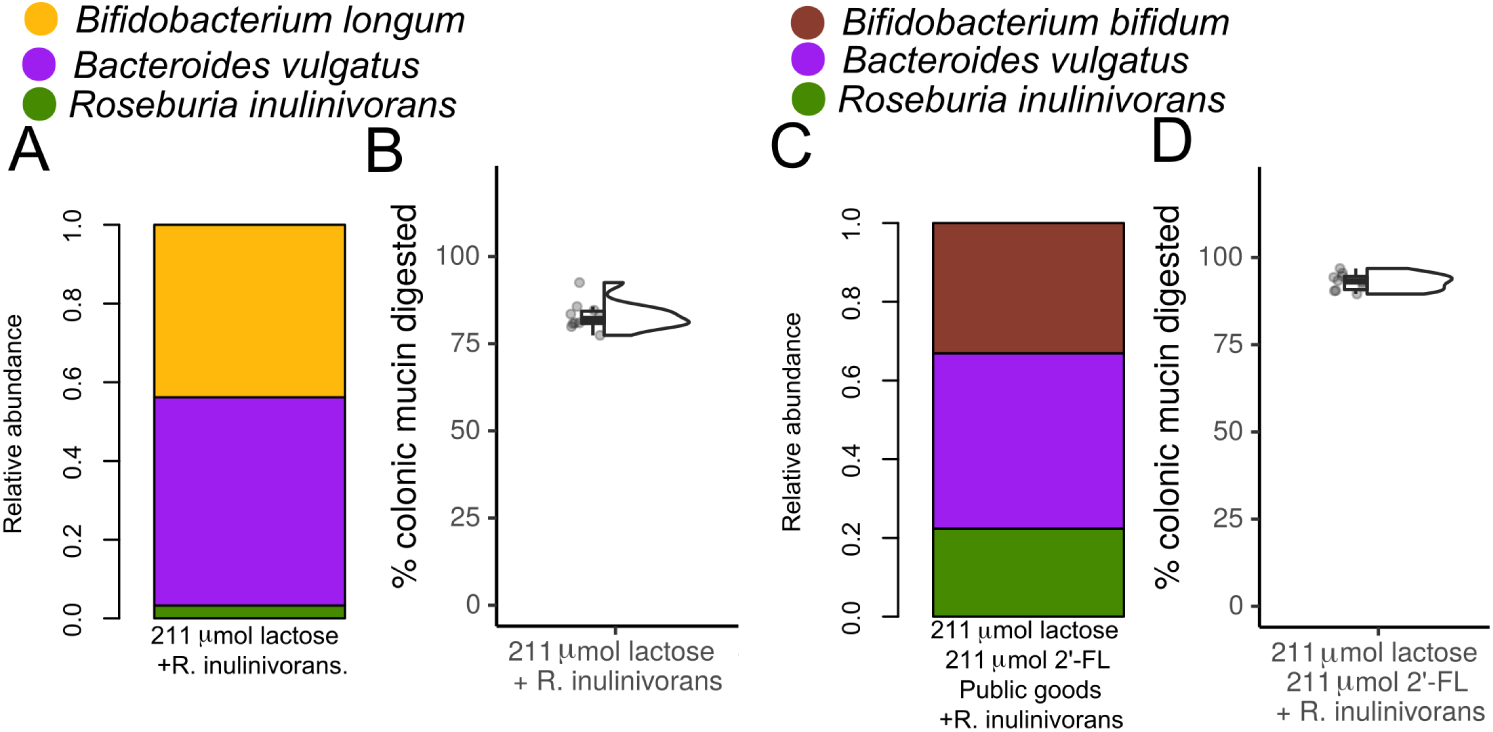
(A) Average relative abundance of *B. longum*, *B. vulgatus*, and *R. inulinivorans* without oligosaccharides at the end of 21 days. n=10 (B) Amount of colonic mucin digested by the microbiota as a percentage of total mucin released into the gut over the final 60 timesteps of the model for A. n=10 (C) Average relative abundance of *B. bifidum*, *B. vulgatus*, and *R. inulinivorans* with 2’-FL and public good metabolism at the end of 21 days. n=10 (D) Amount of colonic mucin digested by the microbiota as a percentage of total mucin released into the gut over the final 60 timesteps of the model for C. n=10

## 6. Contributions

J.M.W.G., and R.M.H.M acquired funding. D.M.V., J.M.W.G., C.W., and R.M.H.M. conceived and planned the simulations. D.M.V. wrote software used for the simulations. D.M.V. performed the simulations and analysed the data. E.L., J.M.W.G., and R.M.H.M contributed to the interpretation of the results. J.M.W.G., and R.M.H.M. supervised the project. D.M.V. drafted the manuscript. D.M.V., E.L., J.M.W.G. and R.M.H.M. revised and edited the manuscript.

## Acknowledgments

This study was financially supported by FrieslandCampina. E.L., and J.M.W.G. are currently employed by FrieslandCampina. This work was performed using the ALICE compute resources provided by Leiden University.

